# Ecdysone Orchestrates Notch and Broad Symphony to Craft Epithelial Cell Shape Change

**DOI:** 10.1101/2023.10.20.563225

**Authors:** Gaurab Ghosh, Sudipta Halder, Aresh Sahu, Mohit Prasad

## Abstract

Morphogenesis in the metazoans relies on cell shape transformations that forms an integral component of organ development, form generation and maintenance of tissue homeostasis. Nevertheless, a comprehensive grasp of how the epithelial morphogenesis is modulated in the metazoans remain still elusive. The Steroid hormones play a pivotal role in morphogenesis spanning several organs including the gonads, urogenital tracts, and mammary glands. Employing the *Drosophila* oogenesis model, we investigated the the role of steroid hormone receptor, Ecdysone receptor (EcR) involvement in transforming anterior epithelial follicle cells (AFCs) from cuboidal to squamous shape. Consistent with the fact that the activity of EcR in the AFCs coincides with the timing of cuboidal-to-squamous shape transition, we found that depletion of EcR function impedes the shape transformation of AFCs. We report that EcR doesn’t impair the follicle cell fate, but impedes the morphological change by restricting the remodelling of lateral and adherens junctions. Employing the classical genetic tools and immnohistochemistry, we show that EcR limits the Notch-Broad axis to facilitate alteration of the shape of AFCs. Our study suggests a mechanistic model where Ecdysone signalling, via the Notch pathway, finetunes the activity of non-muscle myosin heavy chain zipper, prompting AFC shape transition. In sum, our work illuminates how Ecdysone signalling orchestrates epithelial follicle cell morphogenesis during metazoan development.

---

- *Spatio-temporal upregulation of Steroid Hormone cascade induces cell shape change during Drosophila oogenesis*.
- *Fine Tuning of Notch pathway is critical to facilitate cuboidal-to-squamous transition*.
- *Steroid Hormone, Ecdysone, regulates cell shape change by modulating non-myosin II heavy chain, Zipper*.
- *Activity of Zipper is crucial for epithelial morphogenesis*.

## Introduction

Cell shape transition is an important phenomenon that aids several processes including cell movement, organ morphogenesis, and tissue remodelling. How cells change shape is an active area of research primarily because any mis regulation in this process not only results in congenital defects like sickle cell anemia but also stimulates tumor formation.

The epithelial cells are of one of the most abundant cell types present in the complex metazoans. Their versatile shape and form help them to perform diverse function spanning protection, absorption, secretion, sensing the environment and modulating their response to extracellular signals (Larsen et al., 2020). The basic characteristics of epithelial cells are their unique shape, distinctive apical-basal polarity and expression of specific cell junction proteins. The transitioning of epithelial cells from one shape to another is known as epithelial morphogenesis and it plays a critical role in aiding gastrulation, organ development to potentiating tissue repair and regeneration. For example, in developing mammalian eye, transformation of ectodermal cells from cuboidal to columnar fate is critical for forming the functional lens (Zwaan and Hendrix, 1973). In addition, epithelial cell shape transformation is critical for shaping the various organs including the neural tube, tracheal branching, mammary gland development, and limb differentiation (Kass et al., 2007; Macias and Hinck, 2012; Woodward et al., 2001). Given the wide implication of epithelial morphogenesis, there is a constant attempt to identify factors that regulate this process.

Over the years, *Drosophila* oogenesis has emerged as an excellent model to examine various aspects of metazoan epithelial morphogenesis spanning cell fate specification, group cell movement, and cell shape transformations. A female fly has a pair of ovary, each composed of multiple strings of ovarioles. The ovarioles contain egg chambers in a linear progression of maturity, wherein each chamber goes through 14 stages of development to produce a mature egg. An egg chamber is made up of 16 germ cells surrounded by a layer of cuboidal epithelial cells called the somatic follicle cells. As the egg chamber matures from early to mid-stages, the cuboidal follicle cells undergo a dramatic shape transformation (Horne-Badovinac and Bilder, 2005; Wu et al., 2008). In response to Dpp signaling, the most anterior follicle cells (AFCs) flatten to cover half of *Drosophila* egg chamber(Brigaud et al., 2015). The squamous shape flattened cells, referred to as “stretched cells”, eventually cover the germline nurse cells by stage 10 of *Drosophila* oogenesis. The conversion of cuboidal cells to squamous cells involves the shortening of lateral domains and expansion of the apical and basal domains. This shape transition is known to be aided by Notch signaling, serine threonine kinase Tao and growth promoting Target of Rapamycin signaling (Gomez et al., 2012; Grammont, 2007; Halder et al. 2022). Recently, we have shown that germline-soma communication in developing egg chambers too plays a vital role in the aiding the shape change of the AFCs (Sahu et al., 2021). Though we have some information how the AFCs change shape, the identity of factors and interplay of signaling cascades that modulate the acquisition of the squamous epithelial destiny is still far from clear(Gomez et al., 2012; Halder et al., 2022). Intriguingly, Ecdysone, the steroid hormone, is active in the shape transitioning AFCs during mid oogenesis(Jang et al., 2009). Thus, we set out to examine the role of Ecdysone signaling in mediating the shape change of the cuboidal cells to squamous fate.

Ecdysone is known to triggers different stages in the life cycle of flies including embryonic maturation, larval moulting and metamorphosis in the pupae (Chávez Marcela V. et al., 2000; Jiang et al., 1997; Zirin et al., 2013). Since tissue remodelling or tissue removal forms an important aspect of metamorphosis, Ecdysone is known to induce the expression of genes associated with apoptosis and differentiation (Cakouros et al., 2002; Huet François et al., 1995; Jiang et al., 1997). The Ecdysone binds to heterodimeric nuclear hormone receptors, Ecdysone receptor/Ultraspiracle (EcR/USP), to regulate expression of downstream Ecdysone response target genes. Apart from morphogenetic events, Ecdysone pathway also modulates lipid homeostasis, gene amplification and cell migration (Jang et al., 2009; Sieber and Spradling, 2015; Sun et al., 2008).

Here we demonstrate a novel role of Ecdysone signaling in epithelial morphogenesis. By depleting the function of EcR, we show that Ecdysone signaling is required for affecting the shape change of cuboidal cells to squamous fate. By examining various cell fate markers, we conclude that the defect in the cell shape transition in EcR depleted AFCs is not due to altered follicle cell fate specification. Our results suggest that Ecdysone signalling functions through Notch to modulate the acto-myosin contractility in the shape transitioning AFCs. Altogether this study gives new insights on the role of steroid hormones in regulating epithelial morphogenesis, which will not only help us to understand the basics of metazoan development but would have implications in tumor biology too.

## Results

### Ecdysone regulates cuboidal to squamous shape transition in follicle cells

Previtellogenic to vitellogenic transition in *Drosophila* oogenesis is characterized by dramatic increase in the egg chamber size, specification of migratory border cells and conspicuous follicle cell shape transition. Interestingly this phase transition also coincides with the activation of Ecdysone signaling (EcR) in the anterior follicle cells that undergo shape transformation from cuboidal to squamous fate (**Fig. 1C-E),** (Jang et al., 2009). Unlike the expression of Ecdysone receptor in all the follicle cells of developing egg chambers, the Ecdysone Response Element fused to ß-galactosidase gene gets activated in the shape transitioning AFCs *perse* (**Fig. 1A-E)**. This prompted us to examine the role of EcR Signalling in shape transitioning AFCs.

**Figure 1:**
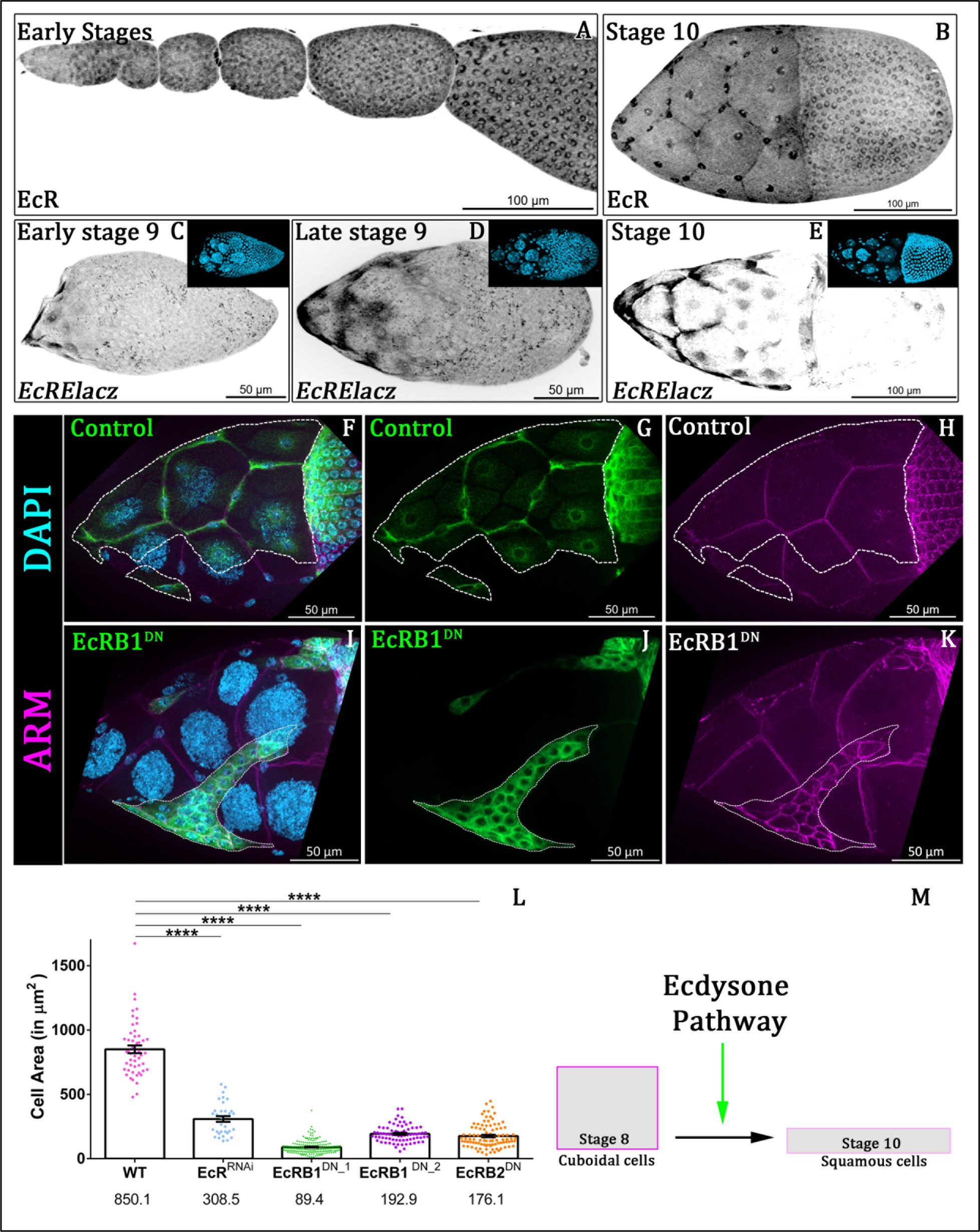
Ecdysone Pathway requires for cuboidal-to-squamous transition. **(A-B)** Distribution of Ecdysone Receptor (EcR) in early Oogenesis**(A)** and at stage 10**(B)** **(C-E)** Stage-wise activation of Ecdysone Pathway in anterior follicle cells. at stage 9**(C-D)** and at stage 10**(E)** **(F-L)** Comparison of Cell Area in Control **(F-H)** and EcRB1^DN^ **(I-K)** expressing anterior follicle cells marked by Flipout Gal4 mediated mCD8GFP expression. **(L)** Graphical representation of cell area in control flipout cells and EcRB1^DN^ expressing Flipout cells. Mean values indicated below each genotype in µm^2^. Armadillo in magenta. **(M)** Schematic representation. Error bars indicate SEM. **** indicates p-value < 0.0001 in Students’ t-test.

To test our hypothesis, we down regulated EcR function in random subset of the AFCs using flipout approach over expressing EcR^RNAi^ construct (TRiP.HMJ22371). The TRiP HMJ22371 construct targets all the isoforms of EcR and over expression of the same resulted in smaller cells. We quantified the size of the EcR RNAi overexpressing cells and found that they are 2.7-fold smaller compared to the nearby control cells. (UAS mCDGFP-850.1±30.16 SEM µm^2^; UAS EcR^RNAi^-308.6±22.02 SEM µm^2^) **(Fig. S1A-C, 1L).** Since the genetic location of EcR precludes any kind of mutant analysis in the adult organism, we perturbed EcR function using the dominant negative (DN) constructs to validate the RNAi results observed above. *Drosophila* genome codes for three EcR isoforms identified as A, B1, and B2, on the basis of different N terminal domains. We used DN constructs for B1 and B2 EcR isoforms independently. The DN constructs targeting various forms of EcR are amino acid alterations that prevent ligand binding, but sequesters the co-receptor, Ultraspiracle thus inhibiting the downstream signaling. Similar to EcR ^RNAi^, clonal over expression of DN Constructs against EcRB1 and EcRB2 isoforms resulted in nine-fold and four-fold drop in the cell area of the AFCs respectively compared to that observed in the control (UAS mCDGFP-850.1±30.16 SEM µm^2^, UAS EcRB1^DN^-89.5±3.4 SEM µm^2^, UAS EcRB2^DN^-192.9±7.6 SEM µm^2^) **(Fig. 1F-L, S1D-F)**. We obtained similar result with another DN construct against EcRB1 isoform (EcRB1^DN^-176.1±9.2 SEM µm^2^) **(Fig. 1L)**. To rule out the possibility of any secondary effect of the FLIPOUT GAL4 construct itself on the shape transitioning follicle cells, we drove EcRB1^DN^ in all the follicle cells by *GR1*-GAL4 driver that expresses in all the follicle cells from early oogenesis. To circumvent the lethality associated with downregulation of EcR in the embryonic stages of *Drosophila* development, the GAL4 activity was suppressed by incubating the cross at 16°C and by including GAL80^ts^, which is a GAL4 repressor. Shifting the progenies to non-permissive temperature inactivated the repression by GAL80^ts^ thus permitting GAL4 expression only in the follicle cells of the developing egg chambers. Satisfyingly, we observed smaller cells at the anterior end similar to that observed when EcR function was downregulated in random subset of AFCs (Control-1081±36.06 SEM µm^2^; EcRB1^DN^ −390.6± SEM µm^2^) **(Fig. S1G-M)**. Altogether from the results above, we conclude that EcR function is required in the AFCs undergoing shape transition from cuboidal to squamous fate **(Fig. 1I)**.

### Perturbation of Ecdysone pathway does not affect the fate of Anterior follicle cell

We know that the follicle cells undergo mitotic to endocycle switch during mid oogenesis (Sun et al., 2008). Since, we observed a large number of ecdysone depleted cells clustered together on the anterior side, we explored the possibility whether EcR knockdown prolonged the mitotic phase resulting in the presence of supernumerary follicle cells. To test this, we immunostained EcR perturbed egg chambers with phosphoHistone H3 (PH3), a marker for mitotically active cells. We found EcR depleted non-flattened clonal cells are PH3 negative indicating that they are not mitotic as speculated **(Fig S2A-C)**. To still rule out the possibility of delayed endocycle onset in the EcR depleted AFCs, we assessed for Number of PH3 in the follicle cells of stage 5 and 6 egg chambers. We did not find a significant difference in PH3 positive cells in EcRB1^DN^ (10.76±7.62 SEM) with respect to control follicle cells (11.07±4.131 SEM, *p=0.2448*) **(Fig. 2A-E).** As no PH3 positive cells were observed after stage 6, we conclude that the defect in the shape changing AFCs is not an outcome of prolonged mitotic phase. The EcR-depleted follicle cells undergo normal mitotic to endocycle transition similar to that observed for the controls (n>40) **(Fig. S2D)**

**Figure 2:**
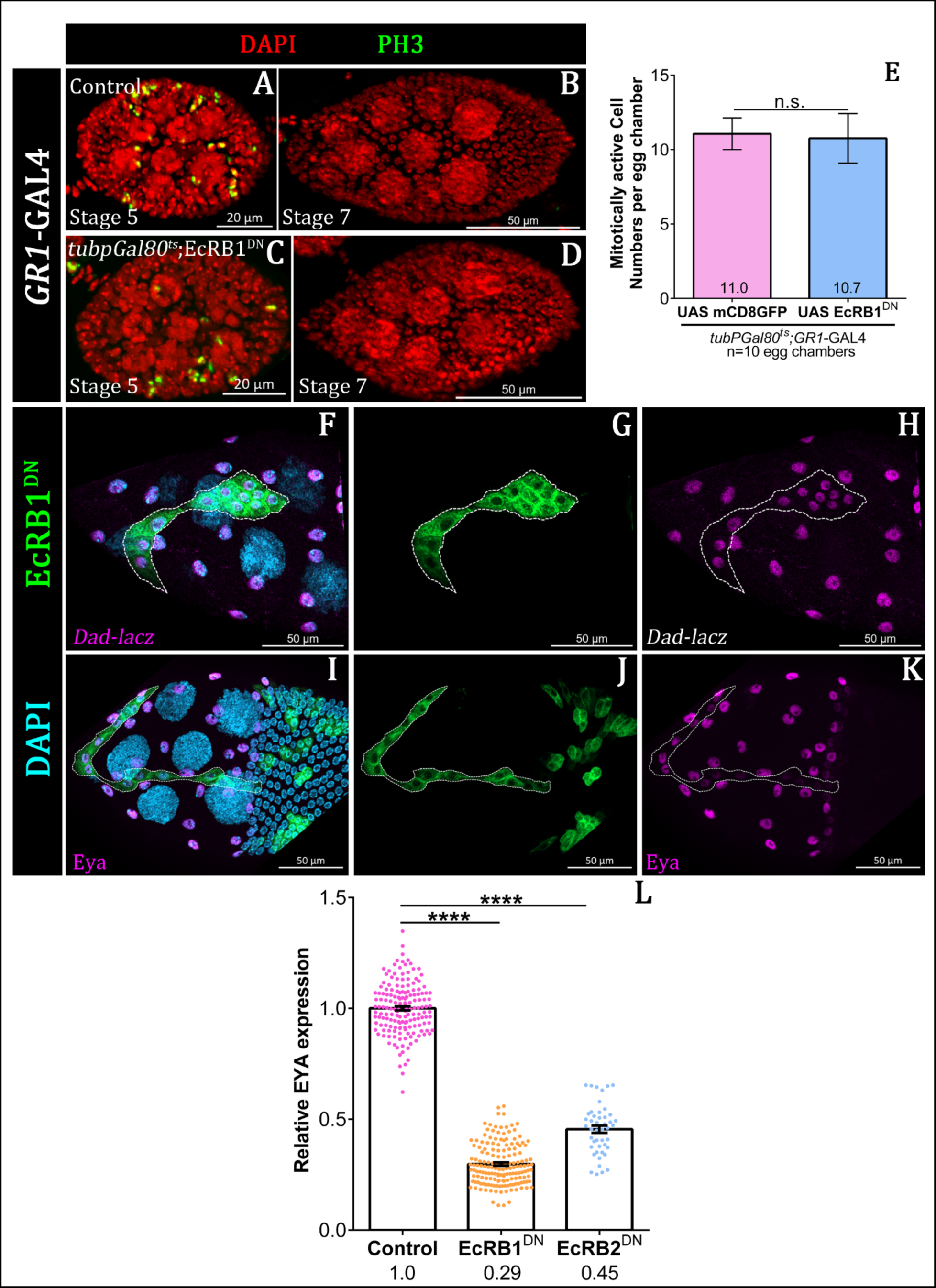
Perturbation of Ecdysone Pathway does not impede Squamous cell Fate. **(A-E)** Comparison and quantification of PH3 staining in GR1-Gal4 driven Control and EcRB1^DN^ follicular epithelium. PH3 count was unaffected between Control and Ecdysone pathway depleted follicle cells. DAPI in red and PH3 in green. **(F-K)** EcRB1^DN^ expressing anterior follicle cells marked by Flipout Gal4 mediated mCD8GFP expression. **(D-F)** Stage 10 egg chambers of indicated genotypes. *dad-lacz* and DAPI imaged in Maganda and cyan respectively. EcRB1^DN^ expressed cells express are *dad-lacz*. **(I-K)** Transcription factor Eya is expressed in anterior follicle cells. Over expression of EcRB1^DN^ by Flipout-Gal4 exhibit lower level of Eya. DAPI and Eya imaged in cyan and Magenta respectively. **(L)** Plot of relative Eya expression in Control, EcRB1^DN^ and EcRB2^DN^. Error bars indicate SEM. **** indicates p-value < 0.0001 in Students’ t-test.

Next, we were curious to examine if this phenotype was an outcome of compromised cell fate specification of the AFCs in the EcR-depleted background. Typically, the germline nurse cells of previtellogenic egg chambers are surrounded by a single layer of mitotically active cuboidal follicle cells. As the developing egg chamber transitions to the vitellogenic phase, the follicle cells undergo conspicuous morphogenetic transformations to acquire different forms and shapes including migratory border cells, stretch follicle cells, centripetal cells and columnar follicle cell enveloping the growing oocyte. Hence, we next examined the fate of the EcR-depleted follicle cells exhibiting shape change defect. To do this, we first assessed the status of Decapentaplegic (Dpp) signalling pathway, which is known to affect flattening by modulating the timing of the morphogenetic process (Brigaud et al., 2015). We examined the status of phosphorylated Mad and *dad-lacz* that are activated in the AFCs undergoing shape change in response to Dpp signaling. However, we didn’t observe any significant difference in the levels of both pMad and *dad*-lacz between the EcR-depleted and the control follicle cells suggesting that absence of Ecdysone signaling doesn’t impair these cells to respond to Dpp signaling **(Fig. S2D-F, 2F-H)**. Next, we assessed the status of another anterior squamous cell fate marker, Eyes absent (Eya). Eya is expressed in all the follicular cells till mid-oogenesis. At Stage 8, Eya expression is limited to anterior follicle cells and later it expands to the centripetal cells of stage 10B egg chambers (Bai and Montell, 2002). We checked the Eya expression pattern in flip-out clones of EcR-B1 ^DN^ and B2^DN^. We observed that Ecdysone-depleted cells are positive for Eya expression, although it is two to three-fold lower compared to nearby wild-type cells **(Fig. 2I-L)** (Control-1.0±0.008 SEM, EcRB1^DN^-0.29±0.007 SEM; EcRB2^DN^ −0.45±0.01 SEM). Though Eya was reduced but as it was expressed in EcR-depleted AFCs we conclude that their cell fate is not significantly altered. To check if lower Eya expression was indeed the cause for the observed shape transformation defect, we overexpressed Eya in the EcR depleted follicle cells **(Fig. S2J)** (EcRB1^DN^; UAS Eya-148.8±4.02 SEM µm^2^; EcRB1^DN^; UAS mCD8GFP-158.2±4.5 SEM µm^2^ *p=0.0514*). Since we didn’t observe any rescue in the reduced size of shape transforming AFCs this suggests that lower levels of Eya may not be the primary cause of the defects observed in cuboidal to squamous transitions in developing egg chambers supporting our earlier conclusion that the fate of the AFCs may not be significantly altered in EcR background. As Dad-lacz and Eya also mark the centripetal cells, we curious to determine if EcR depleted follicle cells exhibiting shape change defect could be the unmigrated centripetal cells itself. The centripetal cells arise from a small subset of AFCs and migrate in between the nurse cells and oocyte the form the dorsal appendages. To test our hypothesis, we evaluated a centripetal cell marker, *BB127-lacz* in EcR depleted AFCs exhibiting shape change defect. Satisfyingly, we didn’t observe any *BB127* positive cells among the AFCs exhibiting smaller size in EcRB1^DN^ background **(Fig. S2G-I)**, suggesting Ecdysone deficient cells were not derivatives of unmigrated cells. Altogether our results above suggest that the defective cuboidal to squamous cell shape phenotype observed in EcR depleted AFCs is not because of any significant alteration in various follicle cell fates.

Next, we set out to investigate as to how the EcR was mediating shape change in the AFCs. As adherens junction remodelling is critical for cuboidal to squamous cell shape transition, we examined the components of adherens junction in EcR depleted follicle cells(Brigaud et al., 2015; Grammont, 2007; Sahu et al., 2021).

### Ecdysone requires for Cell junction remodelling

Epithelial cells are polarized cells with apical, lateral, and basal domains. During cuboidal-to-squamous cell shape transition, the remodelling of adherens junction is important for facilitating the cell shape transition associated with cuboidal-to-squamous fate in the follicular epithelium (Grammont, 2007). DE-Cadherin and Armadillo, important constituents of adherens junction, are abundant in the apical and subapical region of AFCs till stage 8 and are dynamically pulled off from the anterior follicle cells to facilitate cuboidal-to-squamous transition **(Fig S3A-B)**. We next tested the status of adherens junction in the EcR depleted follicle cells. In stage 8 egg chambers, we didn’t find any significant difference in the distribution and the levels of adherens junction components (Armadillo and DE-Cadherin) between the wild type and EcR depleted AFC **(Fig. 3A^1^-A^2^)**. However, from late stage 8 to early stage 9, we observed armadillo accumulation in the apical region of EcR depleted AFCs, while it was undetected in the neighboring control follicle cells. **(Fig. 3B^1^-C^4^)**. In late stage 9 and stage 10 egg chambers, we observed higher levels of DE-Cadherin (DE-Cad) and Armadillo (Arm) in the EcR depleted AFCs compared to the controls (**Fig. 3D^1^-E^3^, S3C-H)**. This observation suggested that disassembly and the timely removal of adherens junctions components were affected in EcR-depleted cells. We noticed same observation in case of E-cadh **(Fig. S3C-H)**.

**Figure 3:**
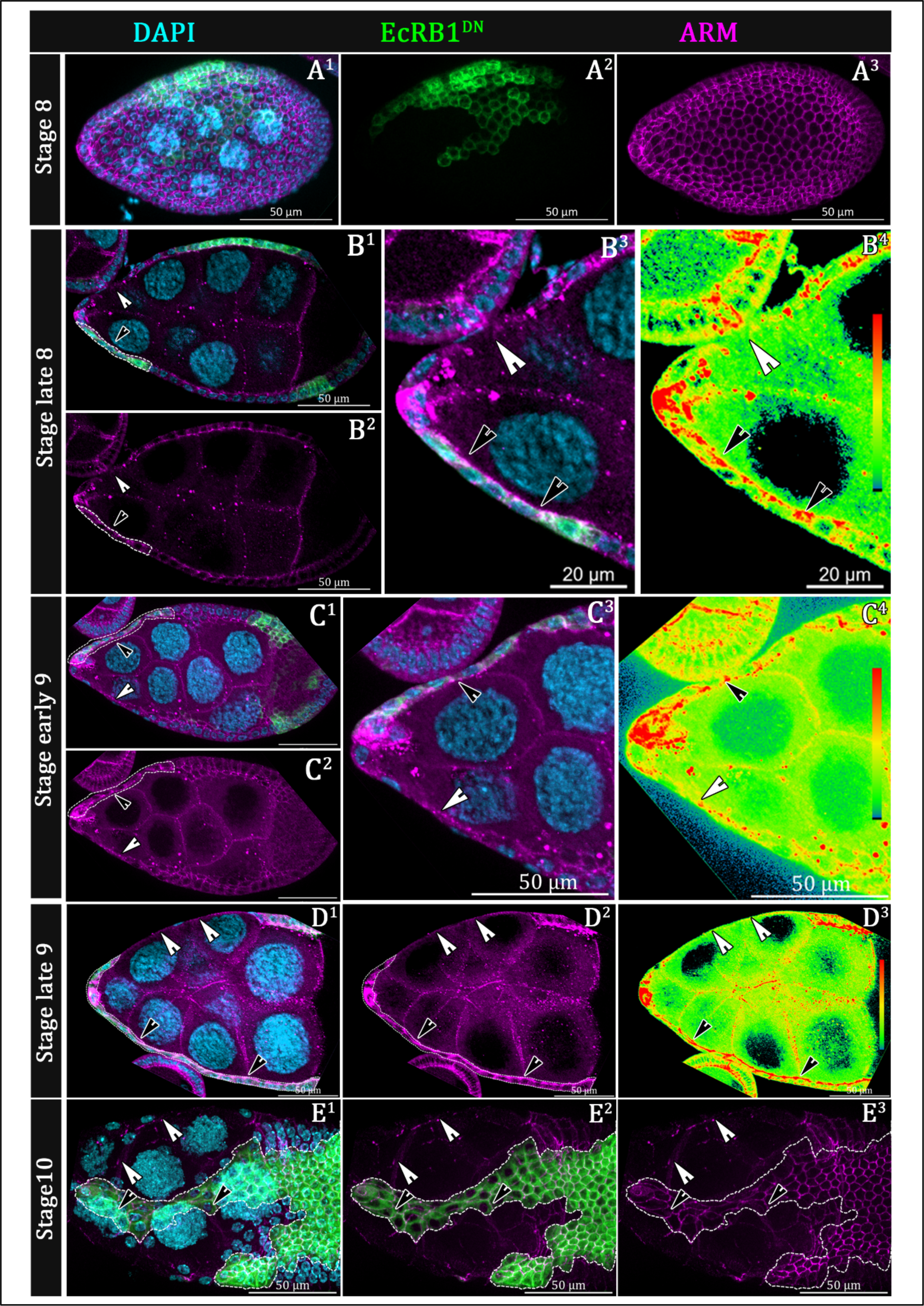
Ecdysone pathway modulates adherens junction to facilitate transition of AFCs. **(A^1^-E^3^)** Stage wise armadillo expression in control and EcRB1^DN^ clonal population. EcRB1^DN^ expressing anterior follicle cells marked by Flipout Gal4 mediated mCD8GFP expression and Armadillo and DAPI in magenta and cyan respectively. **(A^1^-A^3^)** No significant change in distribution of armadillo at stage. EcRB1^DN^ clonal cells are marked by dotted white lines. **(B^1^-E^3^)** Accumulation of armadillo in apical region of Stage late 8 **(B^1^-B^3^),** Stage early 9 **(C^1^-C^3^)**, Stage late 9 **(D^1^-D^3^),** Stage 10 **(E^1^-E^3^)** White arrowheads mark distribution of armadillo in the stretched follicle cells and black arrowheads mark accumulation of armadillo in unstretched follicle cells.

Next we were curious if retention of Arm was primarily responsible for the shape change defect observed in the AFCs. To test this, we genetically removed a copy of Ecad (shotgun) in the EcR background and evaluated the size of the AFCs. Since we did not find any appreciable difference in the size of the EcR-depleted AFCs in the wild type and DE-Cadherin heterozygous background. Neither removal of a copy Arm rescued the smaller size associated with EcR depleted AFCs. Altogether, our results suggest that altered distribution of adherens junction components may not the primary reason for observed cell shape transition phenotypes of AFCs in EcR ^DN^ background (EcRB1^DN^-81.5±3.9 SEM µm^2^; arm^2^; EcRB1^DN^-158.1±5.2 SEM µm^2^, shg^p34-1^, EcRB1^DN^-100.1±5.0 SEM µm^2^) and (EcRB1^DN^; UAS Shg^RNAi^-126.4±2.4 SEM µm^2^; EcRB1^DN^; UAS mCD8GFP-120.4±6.2 SEM µm^2^) **(Fig. S3L, 4H)**.

Next, we wondered if EcR ^DN^ over expressing AFCs, are deficient in removing the lateral protein Fascilin 2? Fasciclin 2 (Fas2) is a cell adhesion molecule from the lateral domains and its timely removal is requisite for transition of AFCs from cuboidal to squamous fate (Gomez et al., 2012; Halder et al.). Generally, Fas2 is expressed till stage 7, after which Fas2 is removed from the lateral membrane of anterior follicle cells (Gomez et al., 2012; Halder et al. 2022).When we generated clones over expressing EcRB1^DN^, we observed retention of Fas2 till stage 9 **(Fig. 3A^1^-A^4^)**. Similarly, we also assessed the distribution of two other lateral membrane components – Discs large (Dlg) and Coracle (Cora) in the EcR depletion background and observed similar retention like Fas2. Unlike Fas2, Dlg and Cora retention phenotype was evident till stage 10 **(Fig. S3F-K)**. Next, we wondered if the retention of lateral domain proteins in background of EcRB1^DN^ was indeed the primary cause of the stretching defect observed.

To address this, we downregulated Fas2 function in the background EcRB1^DN^ and examined the size of the AFCs. Satisfyingly reduction of Fas2 function in the EcRB1^DN^ background gave a partial rescue in cell area when compared EcRB1^DN^ alone (EcRB1^DN^; UAS mCD8GFP-120.4±6.2 SEM µm^2^; EcRB1^DN^; UAS Fas2^RNAi^-368.1±22.68 SEM µm^2^) **(Fig. 4B-H)**. This prompted us to conclude that EcR downregulates Fas2 in the AFCs to enable follicle cell stretching. Interestingly we also observed a reduction of adherens junctional protein, DE-Cadherin in the partially rescued cells within compared to the control (EcRB1^DN^;UAS mCD8GFP)**(Fig. 4G)**.

**Figure 4:**
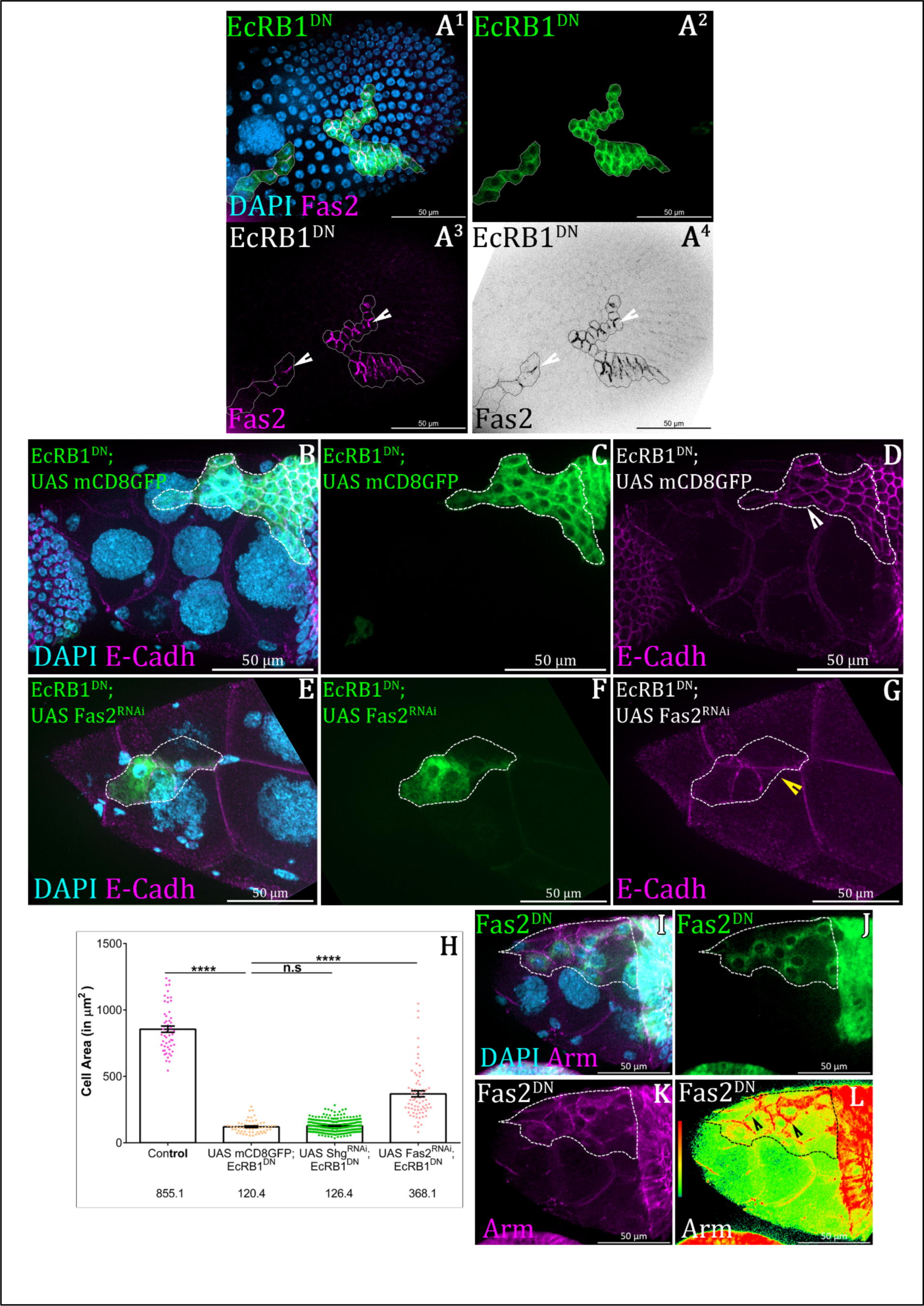
Ecdysone pathway modulates lateral junction to facilitate transition of AFCs. **(A)** Schematic diagram of position of lateral and adherens junction proteins. **(B^1^-B^4^)** Accumulation of Fas2 at Stage late 9 in EcRB1^DN^ clones. DAPI in Cyan, Flipout mediated clones are in Green and Fas2 in Magenta. **(C-H)** Genetic Interaction between EcR and Fas2. Removal of Fas2 partially Green of indicated genotype and E-Cadh in Magenta. **(I)** Quantification of cell area of indicated genotypes. **(J)** Accumulation armadillo in Fas2^DN^ overexpressed cells. DAPI in Cyan, Flipout mediated clones are in Green and Arm in Magenta. Error bars indicate SEM. **** indicates p-value < 0.0001 in Students’ t-test.

Then, we sought to ascertain whether the time-dependent removal of Fas2 during the transition from cuboidal to squamous morphology plays a critical role in the remodeling of adherens junctions or not. To address this, we overexpressed Fas2^DN^ in follicle cells by Flipout GAL4. Fas2^DN^ construct that lacks three amino acids in its C-terminal domain. These three amino acids are essential for its physical interaction with the PDZ domain of Dlg but do not impact the polarity of cells (Szafranski and Goode, 2004). Upon overexpression of Fas2^DN^, we noted a noticeable delay in the removal of the adherens junctional protein at stage 10, Armadillo, within the clonal population. This observation suggests that the timely removal of Fas2 is a pivotal step leading to remodeling of the adherens junctions during cuboidal-to-squamous transition **(Fig.4I-L)**.

Cell shape changes is aided by cell junction remodelling which inturn is influenced by the dynamicity in the activity of non-muscular Myosin-II (Brigaud et al., 2015; Grammont, 2007). We next focussed on the signalling cascades that are known to modulate cell junction components in the shape transitioning AFCs.

### Ecdysone limits Notch Pathway to facilitate stretching of AFCs

Since Notch signalling regulates the adherens junction remodelling in the shape transitioning anterior follicle cells, we investigated its status in the Ecdysone-depleted AFC (Grammont, 2007). Notch receptor is a transmembrane protein with an extracellular domain (ECD) and an intracellular domain (ICD). When activated by the binding of ligand, Delta, it undergoes proteolytic cleavage, releasing the Notch intra-cellular domain (NICD), which translocates to the nucleus and upregulates Notch responsive genes (Bland et al., 2003; Bray, 2006b; Palmer et al., 2014; Sprinzak et al., 2010; Torres et al., 2003) Since analysis of NICD levels and distribution is frequently employed to evaluate the status of Notch signaling, we immunostained EcR depleted with an antibody against the NICD. In the wild type follicle cells, NICD staining was detected both in the membrane and the cytoplasm. However, we observed lower levels of membranous NICD in EcR depleted AFCs compared to that observed in control follicle cells **(Fig. S4G-I).** These findings prompted us to speculate that the absence of NICD from the membrane may indicate hyperactivation of Notch signaling. To test this possibility, we examined the status of Notch pathway using the NRE-eGFP Notch reporter construct. This construct consists of Notch Response Element (NRE) tagged upstream of eGFP, which is activated in response to Notch signaling (Saj et al., 2010). We observed a 3-fold higher intensity of NRE-eGFP reporter in the EcRB1^DN^ overexpressing AFCs compared to the control follicle cells (EcRB1^DN^ −3.1±0.1 SEM; Control-1.0±0.009 SEM) **(Fig. 5A-B)**. Since higher levels of NRE-eGFP was detected in the EcR depleted AFCs, it may suggest that Ecdysone probably functions to restricts the Notch pathway in the shape transitioning AFCs. Interestingly the expression of NRE-eGFP in AFCs continues till stage 8 and tends to decrease in early stage 9 coincident with the activation of *EcRE-LacZ* **(Fig. S4A-F)**.

**Figure 5:**
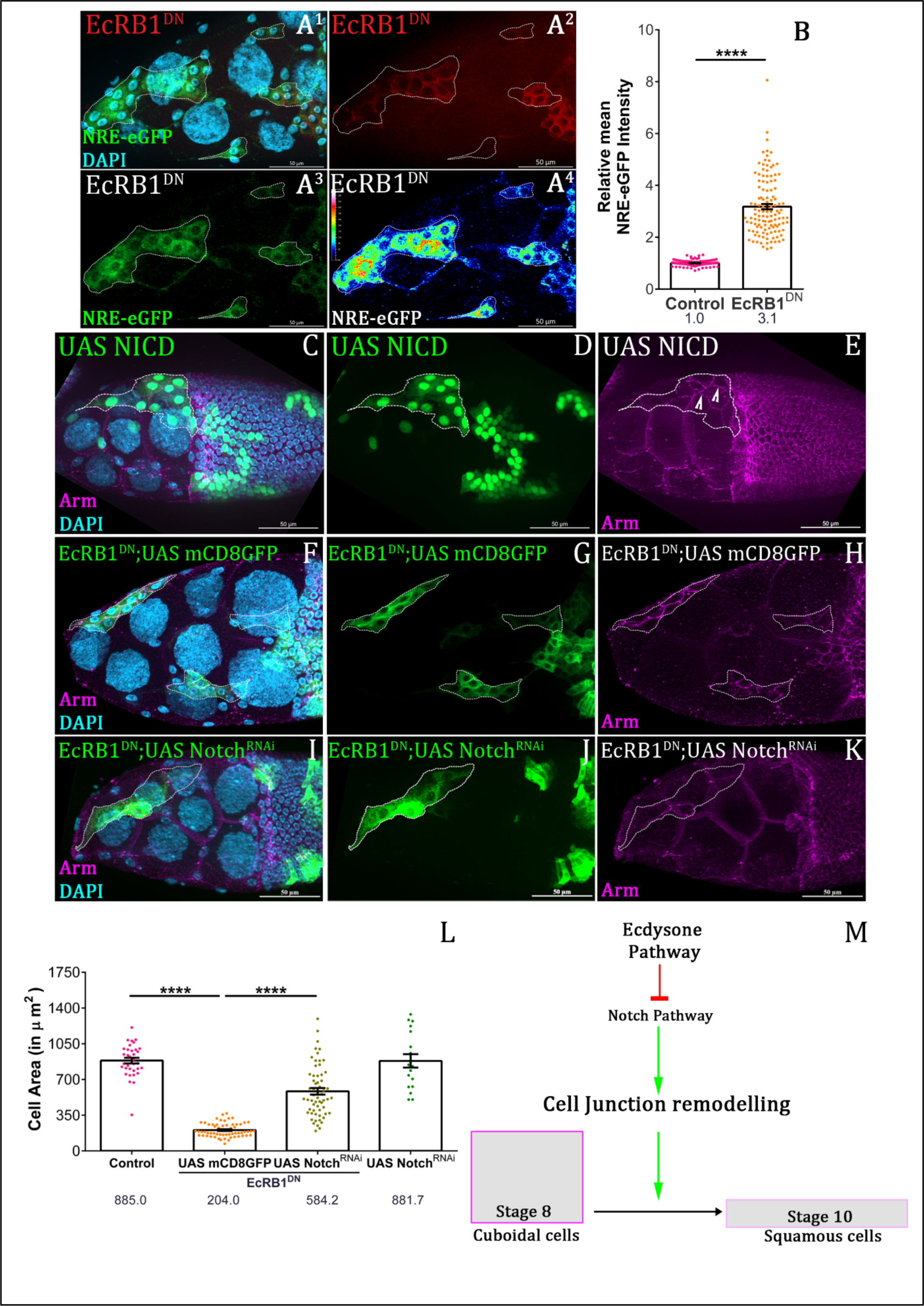
Ecdysone Pathway restricts Notch Pathway to modulate Cell shape change. **(A^1^-A^4^)** Stage 10 egg chamber of indicated genotypes. White dotted line outlines the EcRB1^DN^ expressing unstretched AFCs marked by Flipout mediated myr-RFP. NRE-eGFP expression is in Green, and DAPI in cyan. **(B)** Quantification of the NRE-eGFP of and EcRB1^DN^. **(C-E)** NICD over-expressing unstretched AFCs in AFCs marked by UAS GFP and Armadillo in magenta and DAPI in cyan. **(F-K)** Rescue of stretching defect of EcR-depleted follicle cells by over expression of Notch^RNAi^. Stage 10 egg chambers of indicated genotypes stained with anti-Armadillo in magenta and DAPI in cyan. Clonal cells of indicated genotypes marked by the white dotted lines. **(L)** Quantification of Cell Area of EcRB1^DN^; UAS mCD8GFP, EcRB1^DN^; Notch^RNAi^ and their respective control. **(M)** Schematic Diagram. Error bars indicate SEM. **** indicates p-value < 0.0001 in Students’ t-test.

To test our hypothesis, we next examined if Notch hyperactivation results in morphogenesis defects phenocopying the shape change defected exhibited by EcR AFCs. Agreeably over expression of NiCD resulted in smaller AFCs compared to the control. **(Fig. 5C-E)**. Then, we tried the rescue experiment by downregulating Notch in the background of EcRB1^DN^and evaluated the size of the AFCs. Technically this was difficult to test using standard methods as downregulation of Notch is known to induce excessive proliferation resulting in smaller size follicle cells. So, we standardized the time of heat shock for generating FLIPOUT clone over expressing Notch^RNAi^ so that Notch signaling was down regulated after the follicle cells have undergone normal mitotic to endocycle switch. To check this, we evaluated the level of membranous NiCD along with surface area of the follicle cells. Our premise was to identify a condition where the membranous NiCD is lower but the size of the follicle is not significantly altered **(Fig. S4J-L)**. We found that a heat shock of 45 mins followed by fattening of flies for 16 hours at 29C didn’t not affect the size of the follicle cells appreciably but did result in lowering of membranous NiCD (881.7±65.83 SEM µm^2^) over control (850.1±30.16 SEM µm^2)^ **(Fig. 5L)** (Deng et al., 2001; Sun and Deng, 2005).

We attempted the rescue experiment with EcR^DN^ and NotchRNAi as per the standardized temperature condition mentioned above. Satisfyingly we observed partial rescue in the size of the AFCs when we downregulated Notch in the background of EcRB1^DN^ (584.2±250.3 SEM µm^2^) compared to EcRB1^DN^; UAS mCD8GFP. (204.0±66.36 SEM µm^2^) **(Fig. 5F-L)**.

To further support the above observation, we employed additional approaches to downregulate Notch signaling in background of EcR^DN^. Suppressor of Hairless (*su(H)*) and Fringe (*fng)* are positive regulators of Notch pathway (Bray and Furriols, 2001; Fortini and Artavanis-Tsakonas, 1994; Grammont, 2007; Moloney et al., 2000; Yang et al., 2005). Su(H) is critical for initiating the transcription of Notch response genes with help of other molecules including Mastermind (Mam) and NICD(Bray, 2006b; Bray, 2016; Gomez-Lamarca et al., 2018; Kopan and Ilagan, 2009). While, Fringe modifies Delta ligand aiding Notch activation (Yang et al., 2005). Thus, we introduced *su(H)* and *fng* null alleles as transheterozygotes in the background of EcR depleted follicle cells and evaluate their size. From both of these experiments, we observed partial rescue of the size defect observed for EcRB1^DN^ over expressing follicle cells (EcRB1^DN^-130.2±7.6 SEM µm^2^; EcRB1^DN^; *fng^13^*-370.9±23.05 SEM µm^2^; *su(H)^1^*, EcRB1^DN^ −421.6±44.9 SEM µm^2^) **(Fig. S4M)** (Correia et al., 2003; De Celis et al., 1996). These findings support our premise that Ecdysone modulates the Notch signalling in anterior follicle cells to facilitate the shape change from cubodial to squamous fate.

Overall, our results above suggest that hyperactivity of Notch do is one of the causes of the faulty morphogenesis of anterior follicle cells. Next, we set out to examine as how Notch may modulate the cell shape of the AFCs during *Drosophila* oogenesis.

### Ecdysone pathway negatively modulates Broad-Complex in Anterior Follicle Cells

To extend our findings further, we examined the status of standard downstream targets of Notch in developing follicle cells. Notch signalling is activated in follicle cells during stage 6 egg chambers in developmental egg chambers. Previous work has demonstrated that Notch activation promotes expression of Zinc finger protein Hindsight (Hnt), while suppressing the homeodomain protein Cut (Sun and Deng, 2005). Thus, if conventional Notch pathway is hyper-activated in Ecdysone pathway depleted cells, then we expected alteration in the levels of Hnt and Cut in EcR depleted follicle cells compared to the control follicle cells. Unlike our expectation, we did not observe any alteration in the levels of Hnt and Cut **(Fig S5A-F)**. This observation suggested that an alternate downstream effector is probably regulating morphogenesis downstream of Notch. Coincidently, another downstream target of Notch, Broad-complex, is implicated in modulating mitotic to endocycle transition during *Drosophila* mid oogenesis(Jia et al., 2014).

First, we examined expression of Broad-complex in the background of *EcRE*-Lacz and found that though *EcRElacz* is absent, Broad is present in all the follicle cell up to stage 8 **(Fig. 6A^1^-A^3^)**. During early stage 9 Broad begins to disappear from the anterior follicle cells of developing egg chamber concomitant with activation of *EcRElacz.* **(Fig. 6B^1^-B^3^)**. In stage 10 egg chambers, *EcRElacz* is present only in the flattened AFCs while Broad is absent **(Fig. 6C^1^-C^3^).** The complementary expression pattern of EcRLacz and the Broad complex in the follicle cells led us to speculate if the onset of Ecdysone signalling is essential for Broad downregulation? Satisfyingly, EcRB1^DN^ over expressing clones of follicle cells exhibit Broad retention in the stage 10 egg chambers suggesting that EcR probably negatively modulates Broad expression in the AFCs **(Fig. 7A^1^-A^3^)**. Though Broad is reported downstream target of Notch pathway in stage 5-6 egg chambers, our data suggest this regulation is also operating in the stage 8 egg chambers. Here we revalidated this during stage 8 by checking Broad in the background of Notch^RNAi^ and observed lower levels of Broad in Notch^RNAi^ clones **(Fig. S5G-I)**. Our next question was whether the defective cuboidal-to-squamous transition observed in EcR depleted AFCs is caused by Br upregulation? To address this, first we generated over expressing clones of Br using FLIP-OUT technique. We observed Br overexpressing AFCs were small, retained both Armadillo and Cadherin in stage 10 egg chambers (Control-1069±39.92 SEM µm^2^; UAS Broad Z1-190.9±9.2 SEM µm^2^) **(Fig. 7B-E)**. This defect observed with Br over expression phenocopies the defect observed when EcRB1^DN^ was over expressed in the AFCs (EcRB1^DN^-161.4±4.88 SEM µm^2^) **(Fig. 7E).** However, it is still unclear whether the increase in Broad expression in the background of EcRB1^DN^ is the factor that limits cuboidal-to-squamous shape transition of the AFCs. To test this, we introduced Br null allele, *br^npr-3^*, as a transheterozygote in the background of EcRB1^DN^ and observed a partial rescue of cell area of EcR^DN^ overexpressing anterior follicle cells. (EcRB1^DN^-136.9±9.6 SEM µm^2^; *br^npr-^/+^3^*; EcRB1^DN^-528.6±36.85 SEM µm^2^; Control-850.8±32.63 SEM µm^2^) **(Fig. 7F-L)** This supports our hypothesis that EcR modulates the levels of Br to facilitate the shape change of AFCs.

**Figure 6:**
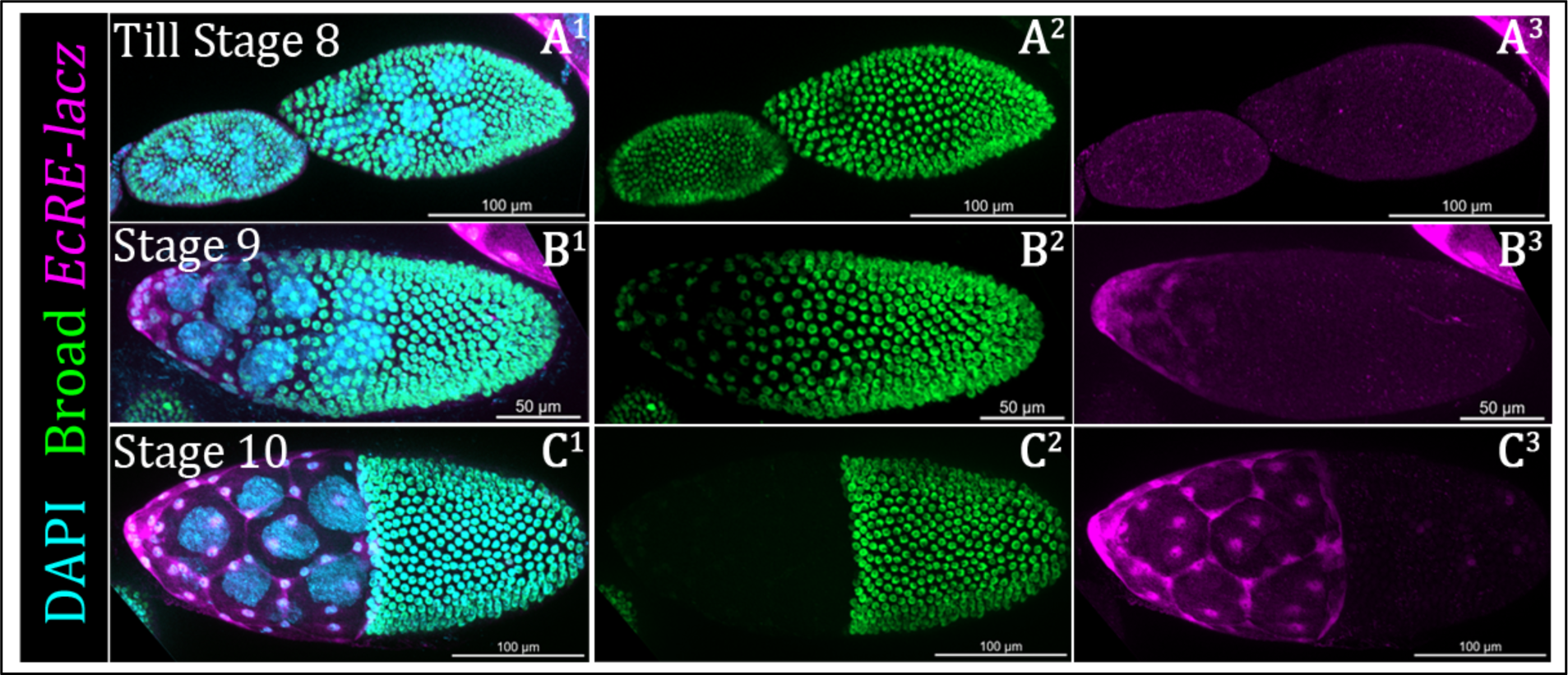
Expression of Broad-Complex and activity of Ecdysone Pathway are mutually exclusive. **(A^1^-C^3^)** Stage wise Expression of Broad in posterior side and *EcRE-lacz* in AFCs at Stage 8 **(A**^1^**-A**^3^), stage 9 **(B**^1^**-B**^3^), stage 10**(C**^1^**-C**^3^). DAPI in cyan Broad in green and *EcRE-lacZ* in magenta.

**Figure 7:**
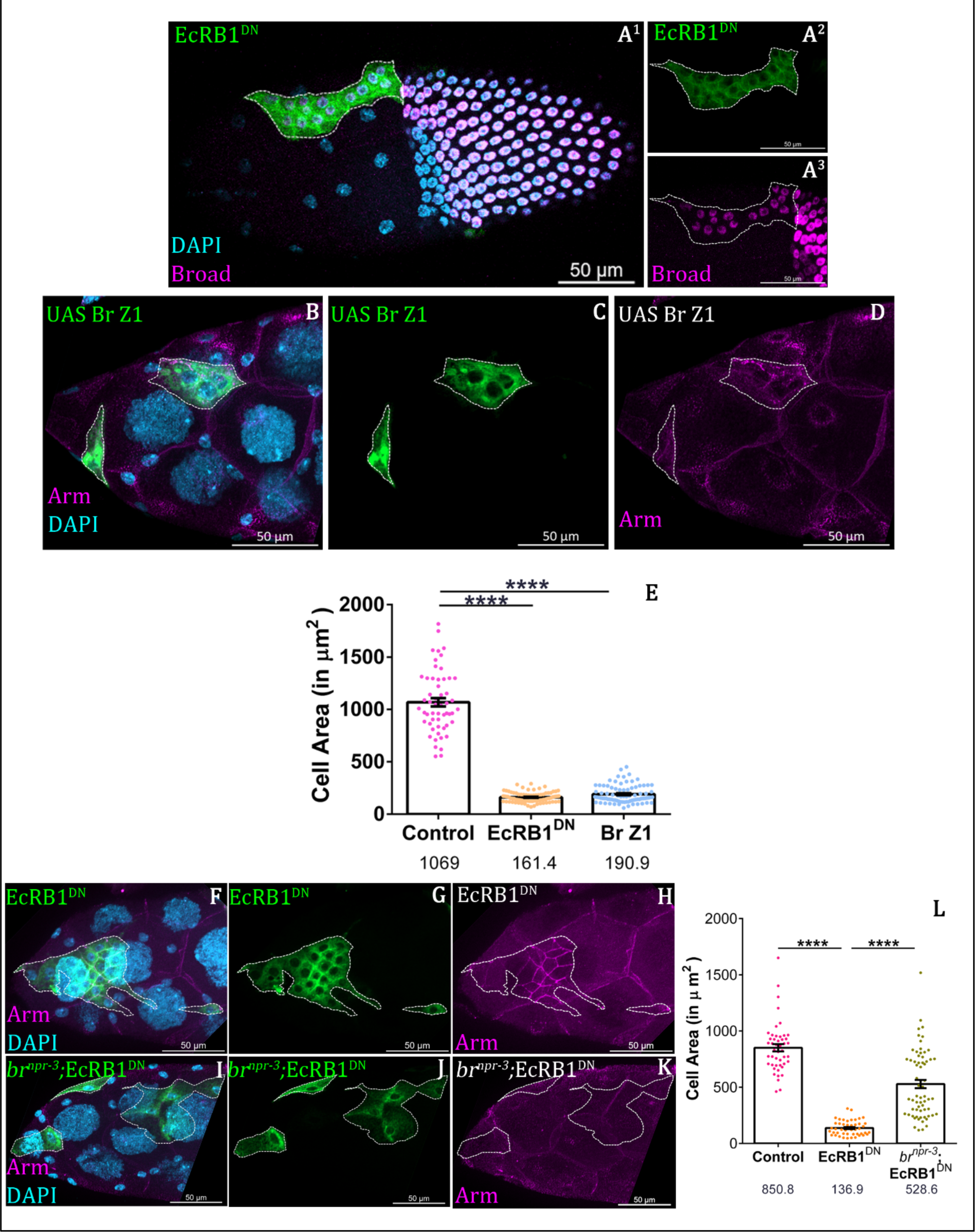
Ecdysone Pathway limits Broad expression. **(A^1^-A^3^)** Broad retains in EcRB1^DN^ expressed anterior follicle cells marked by Flipout Gal4 mediated UAS mCD8GFP. DAPI in cyan and Broad in magenta. **(B-D)** Broad overexpressed Cells in Anterior side exhibit cell shape change defect at stage 10. Cell Area in Control **(F-H)** and EcRB1^DN^ **(I-K)** expressing anterior follicle cells marked by Flipout Gal4 mediated mCD8GFP expression. DAPI in cyan and Armadillo in magenta. **(E)** Quantification of Cell are of Control and UAS Br-Z1 **(F-K)** Genetic removal of Broad (*Br^npr-3^*/+ allele) in EcRB1^DN^ rescues the cell area defects observed upon EcRB1^DN^. Armadillo in magenta and DAPI in cyan. **(L)** Comparison and Plot of cell area across EcRB1^DN^ and *br^npr-3^/+*; EcRB1^DN^. Error bars indicate SEM. **** indicates p-value < 0.0001 in Students’ t-test.

Overall, our findings imply that Ecdysone pathway fine tunes Notch to limit Broad in the shape transitioning AFCs. So far, we have obtained evidence as to how Ecdysone regulates cuboidal to squamous cell shape transition through Notch and Broad. However, identity of the molecule that functions downstream of Notch and can directly modulate junctional protein remodelling is unclear. Hence, we next set to identify a molecule below Notch-Broad pathway that can affect the apico-lateral junction remodelling.

### Perturbation of Ecdysone affects removal of acto-myosin contractility network

Cell shape changes is mediated by in dynamicity of actin-cytoskeleton network that depends on the activity of non-muscular Myosin-II (Brigaud et al., 2015; Grammont, 2007). To dissect out the other possibilities behind the defect, we were inclined to check the status of acto-myosin contractility network in EcRB1^DN^ clonal population. Myosin is required for epithelial remodelling and movement during tissue formation, organogenesis and gastrulation (Gorfinkiel and Blanchard, 2011; Majumder et al., 2012; Skoglund et al., 2008). Non muscle Myo-II is composed of three subunit-*Drosophila* heavy chain is Zipper (Zip), Essential Myosin light chain cytoplasmic (Mlc-C) and Regulatory light chain (MRLC) or Spaghetti squash(sqh) (Karess et al., 1991; Mansfield et al., 1996). Earlier study demonstrated that Zipper and Sqh are critical for aiding cell shape change in the elongating germband in the *Drosophila* embryos (Bertet et al., 2004). During cuboidal-to-squamous transition in anterior follicle cells, Zip and actin accumulates temporally at the junctions and helps in junctional remodelling (Grammont, 2007). Thus, cell shape alterations are also dependent on actin rearrangement in a wide range of developmental contexts. We analyzed the location of Zipper and actin in Ecdysone pathway-depleted follicular cells (Flip out overexpressing clones) to investigate the potential involvement of cytoskeletal alterations. In Stage 10 egg chambers, we found higher levels of Zipper staining in Ecdysone-depleted follicle cells unlike in nearby stretched follicle cells, but we observed lower levels of F-actin in the clonal population **(Fig. 8A-B, S6A-C)**. From the observation we can infer that acto-myosin contractility network is altered in Ecdysone depleted cells. This accumulation of zipper is quite similar with inx2 depleted cells (Sahu et al., 2021).

**Figure 8:**
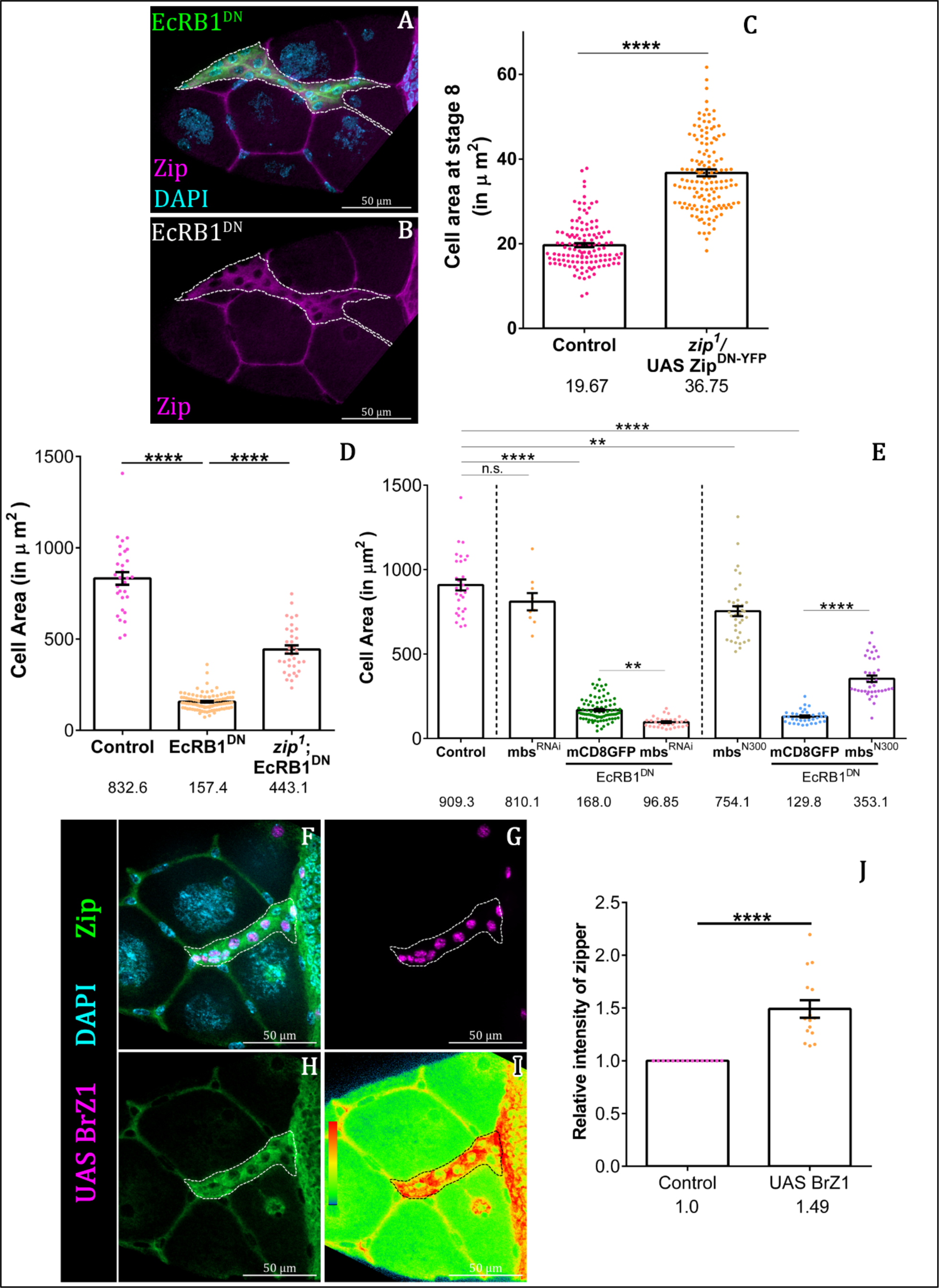
Ecdysone Pathway influences activity of Zipper to accomplish the shape change. **(A-B)** Elevated levels of Zipper observed in EcR perturbed follicle cells compared to the neighbouring control cells undergoing shape change. EcR perturbed cells are marked by mCD8GFP. White dotted outline marks the clonal population, DAPI in Cyan. Anti-Zipper staining is imaged in magenta. **(C)** Quantification of the follicle cell area of indicated genotypes. Zipper activity downregulation gave increased cell size. **(D)**. Quantification of cell area for indicated genotypes. *Zip^1^* trans heterozygotes background rescue the EcRB1^DN^ defect. **(E)** Quantification of cell area of indicated genotypes. Genetic interaction between EcR and mbs. **(F-I)** Elevated zipper expression in the Broad Overexpressed cells. Clonal cells are marked by StingerNLS in magenta, DAPI in cyan and zipper in Green. **(J)** Quantification of relative level of zipper of indicated genotypes at stage 8. Error bars indicate SEM. **** indicates, ns = not significant p-value < 0.0001 in Students’ t-test.

Next, we tested if the Zipper function is instrumental in aiding the shape transition of cuboidal follicle cells to squamous fate? To test our hypothesis, we overexpressed Zip^DN-YFP^ in the follicle cells and examined their size. The construct Zip^DN-YFP^ drives the zip sequences where the actin binding motor domain is replaced by YFP moiety rendering the protein dominant negative. To increase the potency of the Zip^DN-YFP^ effect, this experiment was carried in the background of amorphic allele of *zip^1^*(Dawes-Hoang et al., 2005). Pleasantly, we observed precocious flattening of follicle cells in the stage 8 egg chambers (*zip^1^*/UAS Zip ^DN-YFP^-36.75±0.9 SEM µm^2^; Control-19.6±0.4 SEM µm^2^) **(Fig. 8C Fig. S6E-H).** This suggests that Zipper modulates the shape changes of the transitioning AFCs.

Subsequently we wanted to check if EcR limits zipper in the shape transitioning AFCs? To check this, we sought to rescue the aberrant cell shape change defect observed in EcRB1^DN^ background by reducing zipper levels?. As per our expectation, we observed partial rescue in the cell size of EcR-depleted AFCs by introducing the amorphic allele, *zip^1^*/+ (transheterozygote) in EcRB1^DN^ background (EcRB1^DN^-151.6±17.09 SEM µm^2^; *zip^1^*, EcRB1^DN^-443.1±22.2 SEM µm^2^) **(Fig. 8D)** supporting our hypothesis that EcR indeed limit Zipper function to modulate shape change of AFCs. Having determined that Zip functions downstream of EcR, we were curious if the accumulation of zipper protein *perse* or the activity of myosin protein is impeding the transition from a cuboidal to a squamous cell? To examine this, we overexpressed Zipper (UAS-Zip:YFP) in the AFCs to check if this caused any shape change defect. Unlike our expectation, increase in the overall protein content of Zipper did not exert any discernible impact on the remodelling of adherens junctions nor did it cause any shape change defect of AFCs in stage 10 egg chambers when compared to the control condition (UAS Zip^YFP^-872.9±34.6 SEM µm^2^; Control-907.7±36.2 SEM µm^2^) **(Fig. S6D).**

Subsequently, our inquiry led us to investigate the factors that can exert control over the activity of zipper during epithelial morphogenesis. During dorsal closure in Drosophila embryo, zipper becomes activated through the phosphorylation of its regulatory myosin chain. In this context, we tested whether phosphorylation of the myosin regulatory chain plays a pivotal role in the context of EcR mediated cell shape transition of AFCs. To manipulate the activity of myosin regulatory chain in cuboidal-to-squamous transition, we altered the levels of Myosin binding Subunit (Mbs) within the EcRB1^DN^ genetic context. The Mbs gene is responsible for encoding the regulatory subunit of myosin phosphatase, a protein complex involved in the regulation of myosin activity (Grassie et al., 2011; Kimura et al., 1996). Mbs negatively modulates the catalytic activity of myosin by dephosphorylating myosin regulatory subunit (MRLC) (Tomoaki Mizuno et al., 2002). In order to assess the genetic interplay between EcR and Mbs, we introduced the RNA interference (RNAi) construct targeting Mbs in the genetic background of EcRB1^DN^ over expressing follicle cells. Based on the analysis of the preceding outcome, we can hypothesize that augmenting the myosin activity in the background of EcRDN would lead to an exacerbation of the reduced cell size. This is indeed what we observed when we coexpressed MbsRNAi along with EcR^DN^ in the AFCs (EcRB1^DN^; UAS mCD8GFP-168.0±7.1 SEM µm^2^; EcRB1^DN^;UAS mbs^RNAi^-96.8±5.4 SEM µm^2^) **(Fig. 8E).** On the other hand, we sought to counter the EcRB1^DN^ defect by employing Mbs^N300^construct, which is known to constitutively dephosphorylate MRLC. The rationale behind these interventions was manipulating the activity of myosin-II that could potentially compensate for the defects associated with EcRB1^DN^. Remarkably, our efforts yielded promising outcomes, as we observed a partial of rescue of the cell size in the background of EcRB1^DN^ (EcRB1^DN^; UAS mCD8GFP-129.8±6.11 SEM µm^2^; EcRB1^DN^; UAS mbs^N300^-353.7±18.7 SEM µm^2^) **(Fig. 8E, S6K-N)**

Next, we asked whether hyper activation of Notch pathway phenocopies the zipper accumulation. Pleasantly, overexpression of Broad resulted in elevated levels of Zipper in stage 10, akin to the effect observed in cells overexpressing EcRB1^DN^ **(Fig. 8J-M).** We quantified the Zipper level in stage 8 as all cells are cuboidal in nature and we observed almost 1.5-fold increase of Zipper in BrZ1 overexpressed cells compared to neighboring control cells. **(Fig. 8J, S6O-R).** Together these data suggest that Ecdysone pathway modulates myosin activity to facilitate the stretching of the AFCs.

Our study aimed to elucidate the molecular underpinnings of the phenotypic manifestations resulting from the reduction of the Ecdysone pathway during *Drosophila* oogenesis. The study found that depletion of the EcR leads to increased levels of the Notch Pathway which adversely affect the cell shape transformation of AFCs.. This is probably because the Notch retains the Myosin II activity in the follicle cells thus preventing any shape transition to squamous fate.

## Discussion

Epithelial morphogenesis is critical for acquisition of specific cell shapes during development and tissue repair. Though this process plays a widespread role in form generation and structure maintenance among the metazoans, the underlying molecular mechanism governing this process is far from clear. In this study, we investigated how the follicle epithelial cells transition from cuboidal to squamous fate in the developing fly egg chambers. We found that Steroid hormone, Ecdysone, modulates the shape change of cuboidal follicle cells to squamous fate during mid oogenesis. Given the steroid hormones regulate tissue morphogenesis and organ remodelling in both vertebrates and invertebrates, our results give unique molecular insight into how they would regulate epithelial cell shape transformation in the metazoans.

Our results suggest that EcR restricts Notch signaling in the AFCs in developing stage 8-10 egg chambers. We propose that down regulation of Notch signaling by EcR, depletes the transcription factor Broad and down modulates the Zipper (Myosin II) in the shape transitioning AFCs. Similar to previous reports, we too find that the zinc transcription factor Broad-Complex (BR-C) functions downstream of EcR, however the mode of regulation is indirect (DiBello et al., 1991; Fletcher and Thummel, 1995). Unlike the EcR directly activating the Br-C in the salivary glands, we observe EcR negatively regulates Br-C through Notch signaling in the stage 9 AFCs (Buszczak and Segraves, 2000; Huet François et al., 1995; Thummel CS, 1996). Since Broad is a downstream target of Notch in the follicle cells and its over expression also results in smaller AFCs, we believe that modulation of Zipper activity downstream of Broad is critical for potentiating shape change in the anterior follicle cells. The question remains how Broad is regulating Zipper activity. It would be worth examining the known downstream targets of Broad and check if they do modulate the activity of the Zipper in developing follicle cells.

The next question is how Zip activity is mediating the shape change of the AFCs. It has been reported Zipper along with Diaphanous control E-cadherin endocytosis to assist intercalating epithelial cells during germ band extension in early embryos (Kong et al., 2017; Levayer and Lecuit, 2013). Though we do observe higher levels of DE-Cadherin in the EcR depleted follicle cells, we believe this may not be the primary cause of the shape change defect observed in AFCs. Our speculation stems from the fact that unlike Fasciclin 2 (Fas2) reduction, depletion of DE-Cadherin failed to rescue the smaller size of the EcR depleted AFCs. Fas2 is a member of transmembrane immunoglobulin superfamily that localizes to the lateral membrane of the follicle cells in previtellogenic egg chambers. Interestingly the partially rescued EcR depleted AFCs that over express Fas2 RNAi also exhibit lower levels of DE-Cadherin **(Fig. 4C).** This suggests that accumulation of DE-Cadherin observed in EcR-depleted AFCs may be due to retention of Fas2 itself. Given that Fas2 removal from the lateral membrane is one of the critical steps during transitioning of AFCs from cuboidal to squamous fate, we think that elevated myosin activity in the EcR-depleted AFCs may be resisting Fas2 exclusion thus delaying the shape change from cuboidal to squamous fate. Lowering of myosin activity aids in Fas2 removal, thus aiding the shape of the AFCs during epithelial morphogenesis.

We are still intrigued by the fact that how lower Zipper activity assists in Fas2 removal facilitating stretching and flattening of AFCs? It will be worth examining if Zipper activity *perse* can regulate endocytosis? Given that Zip physically associates with endocytosis regulators Rab6 and Rab7, it would be interesting to test if these molecules regulate the shape change of the AFCs during mid oogenesis (Gillingham et al., 2014).

Our results not only give us novel insight into the Ecdysone pathway and its target in morphogenesis but sheds light on context dependent gene regulation like Broad. Given that hormonal signaling is intricately linked to transcriptional regulation and cell fate transition, our study unveils a previously unrecognized facet of the Ecdysone pathway in mediating cell change in epithelial morphogenesis. In future, how Ecdysone inhibits Notch signaling and Broad-Complex modulates Zipper activity in AFCs is an exciting research question to pursue.

From a broader perspective, our research underscores the significance of precise control and coordination in cell shape change. It implies that fine-tuning the activity of myosin II and conserved signaling pathways is a critical aspect of metazoan development. We believe that our findings will improve our understanding as to how an organism grows, develops, and adapts to their environments by sensing hormonal cues.

Steroid hormones are one of the major modulators of epithelial morphogenesis in metazoans. We know that Estrogen regulates alveolar branching in the developing lungs while Progesterone facilitates ductal elongation in lactating mammary glands (Kass et al., 2007; Woodward et al., 2001). Intriguingly, imbalances of steroid hormones are linked to Breast Cancer development and progression. As tumor formation is associated with aberrant cell shape transitions, it would be interesting to test our findings in higher systems. We believe this may give us an opportunity to identify potential therapeutics to alleviate both diseased conditions and development disorders associated with aberrant cell transformations in the metazoans.

## Material and Methods

### Fly Strains

All stocks and crosses were maintained at 25°C. Experiments involving the expression of RNAi and overexpression constructs of transgenes were fattened at 29 °C for 16 hours., Induction of transgenes using Flipout/AyGal4 was induced by heat shocking at 37 °C 2 times for two day followed by dissection after 2 days. For Notch^RNAi^ experiment, flies were pre-fattened with yeast at 25°C for 24 hours. After that, the clone was induced by heat shock at 37°C for a time of 45mins. After the heat shock, we shifted those flies to 29°C for 16 hours before dissection.

Flies used for this work UAS EcRB1^DN^ (BL-6872,6869), UAS EcR^RNAi^ (BL-52860), UAS EcRB2^DN^ (BL-9450), NRE-eGFP, UAS Br-Z1 (BL-51190), *zip^1^*(BL-4199), *br^npr-3^*, UAS NICD, *EcRE-lacz*, UAS Notch^RNAi^ (BL-28731), *arm^2^,* UAS Fas2^RNai^, UAS Fas2^DN^, UAS shg^RNAi^(BL-32904), *cora^5^*, *shg^p34-1^*(Gift from Dr. Richa Rikhy), *fng^13^*(BL-8552)*, Su(H)^1^*(BL-417), UAS Zip^DN-YFP^(From Dr. Krishanu Ray), UAS Zip^YFP^, UAS mCD8GFP, *dpplacz, dadlacz, BB127lacz* (Gift from Trudi Supbach). The flies were procured from Bloomington *Drosophila* Stock Center (BDSC).

### Immunostaining

Ovaries of female flies were dissected in Schneider’s medium supplemented with 10% fetal bovine serum. Flies were fixed in 4% paraformaldehyde solution (Sigma) for 15-20 mins. Blocking solution was made using 5% bovine serum albumin, 0.3% Triton-X-100 in 1X phosphate-buffered saline (Sigma). For NICD staining we used 0.1% Triton-X-100 in 1X phosphate-buffered saline with 5% bovine serum albumin as blocking solution. Staining was performed using standard protocol.(Saha et al., 2023) Primary antibodies were used at the following concentrations: mouse anti-Armadillo (N2 7A1 at 1:120; Developmental Studies Hybridoma Bank, DSHB), rat anti-DEcadh(1:50; DSHB), mouse anti-coracle(1:50; DSHB), mouse anti-Fas2(1D4 at 1:25;DSHB), mouse anti-Eya(10H6, 1:100;DSHB),mouse anti-Broad Complex(1:50; DSHB), mouse anti-EcR(1:25; DSHB), mouse anti-αTubulin(1:400; Sigma), rabbit anti-β-galactosidase (1:100; DSHB), rabbit anti-GFP (A11122 at 1:500; Thermo Fisher Scientific).

### Intensity measurement

For measuring the NRE-GFP levels, the cloned cells of stage 10 egg chambers were imaged keeping identical exposure time (GFP channel-200-400 ms) and other imaging parameters. The z-planes were merged together to obtain a 2D image. Then, we outlined the ROI by free hands by checking the reporter (myr RFP) and measured the mean GFP intensity. For Control we outlined the same area as experiment in the neighbouring cells and measured mean GFP intensity. The relative mean GFP intensity of experimental follicle cells were plotted as fold change where we kept control as 1 with statistical analysis.Images were acquired in Zeiss Axio observer 7 with Apotome.2 module and analysed with Zen 2012 software.

### Area Calculation

The determination of cell area was performed by delineating the perimeters of individual cells using the Zen Blue software. The determination of the cellular boundary was assessed by evaluating the expression of the GFP marker within the clones, along with the co-staining of armadillo or tubulin.

### Statistical tests

Difference of means determined by Students’ t-test with Mann-Whitney in GraphPad Prism 6.0. All error bars indicate Standard Error of Mean. All figures utilize, the following range for assigning significance of means: p-value *p >0.01* is designated as * and *>0.05* as ns or not significant.

---

**Figure S1:**
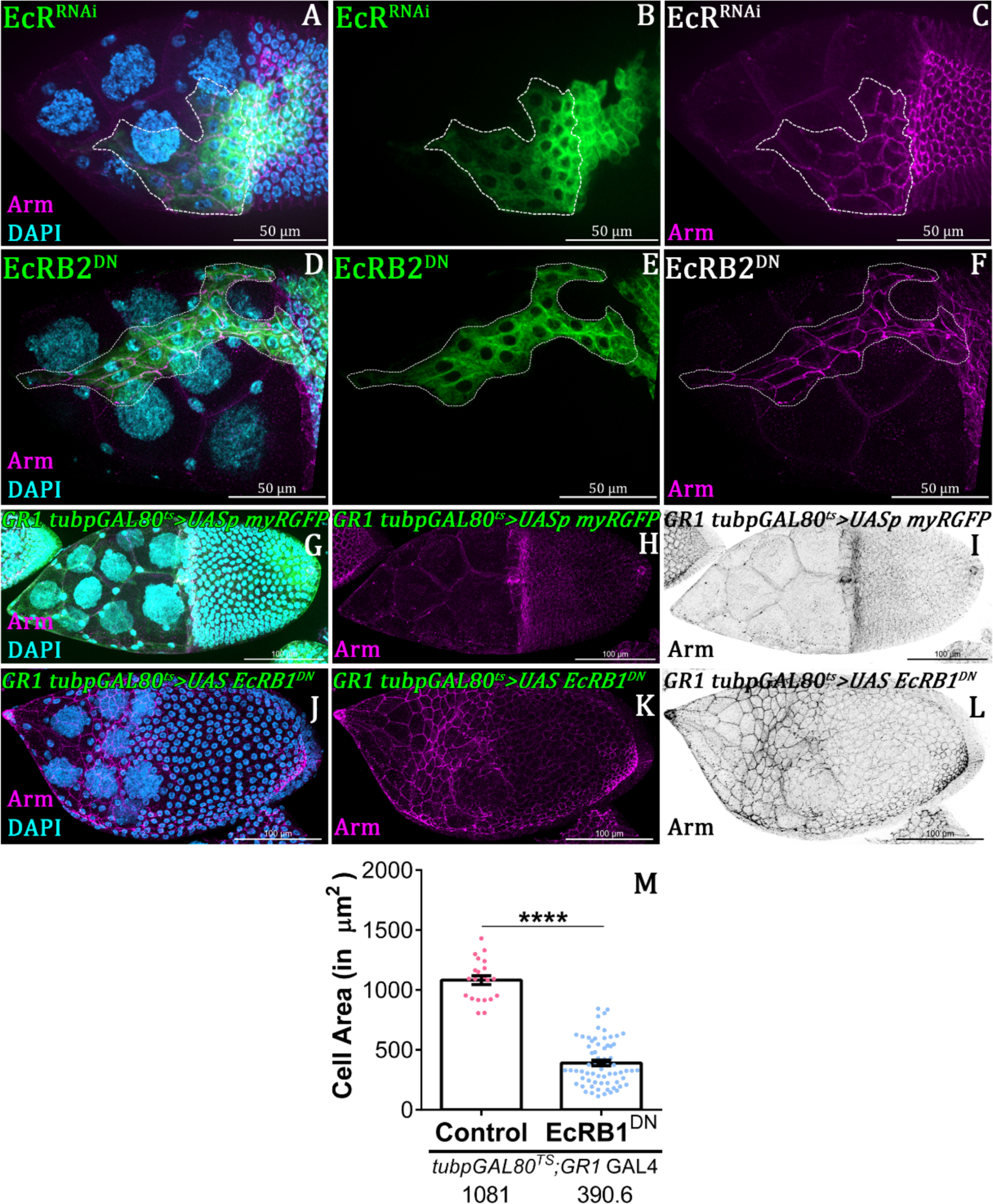
EcR mediates the stretching of the anterior follicle. **(A-F)** Clonal Overexpression of EcR^RNAi^ and EcRB2^DN^ marked by Flipout mediated UAS mCD8GFP. Both gave defect in cuboidal-to-squamous transition. DAPI in cyan and Armadillo in magenta. **(G-L)** Comparison of Cell Area in Control **(G-I)** and EcRB1^DN^ **(J-L)** in background of follicle specific *GR1*-Gal4 driver. DAPI in cyan and Armadillo in magenta. **(M)** Quantification of cell area in the background of indicated genotype. Error bars indicate SEM. **** indicates p-value < 0.0001 in Students’ t-test.

**Figure S2:**
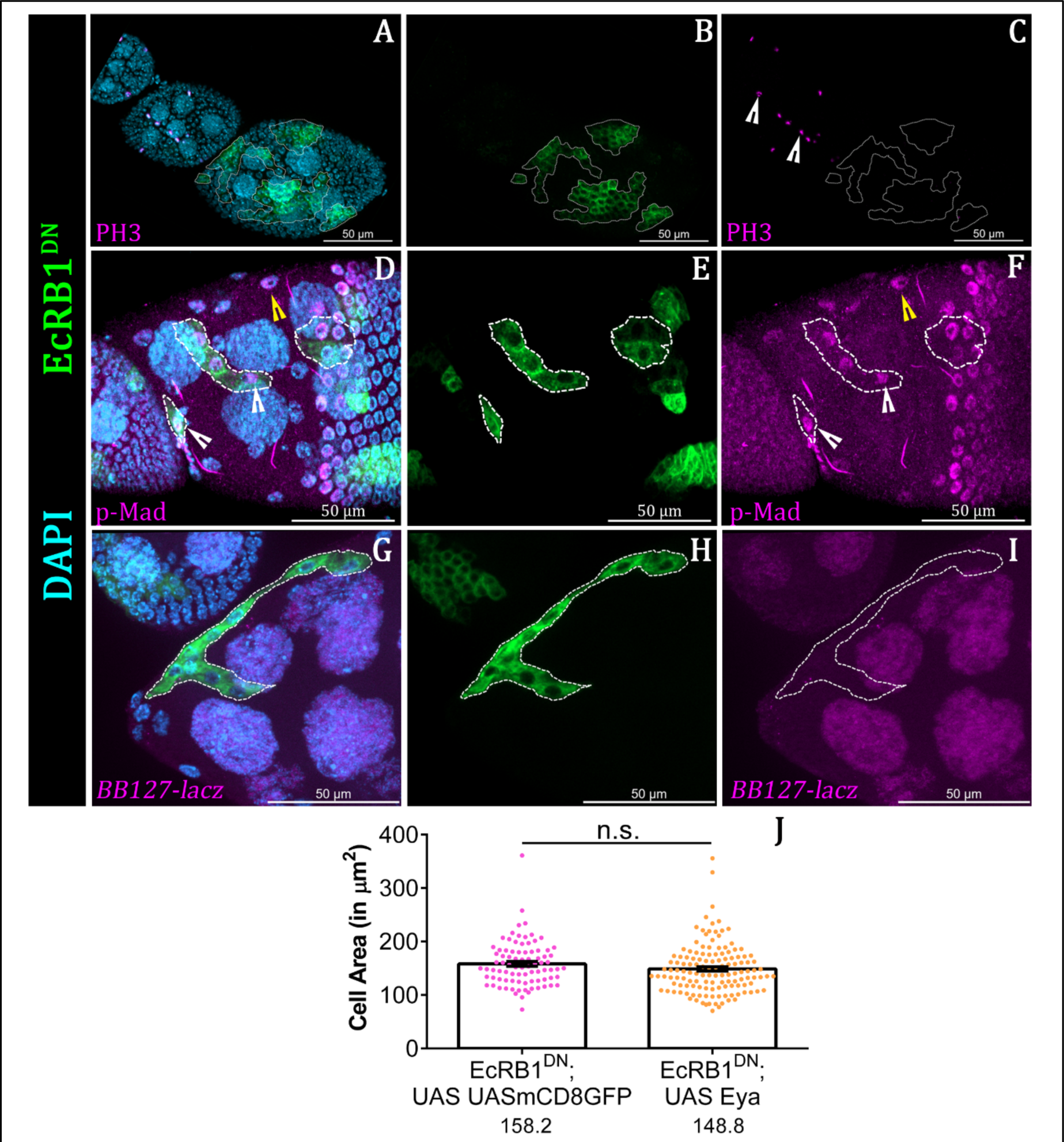
Perturbation of EcR does not affect the fate of AFCs. **(A-C)** Ecdysone Pathway depleted cells are PH3 negative.(white arrow head indicates PH3 positive cells in early eggchambers. Clonal cells are marked by GFP, DAPI in cyan and PH3 in magenta. **(D-F)** EcRB1^DN^ overexpressed follicle cells are positive for pMad staining. Clonal cells are marked by GFP, DAPI in cyan and pMad in magenta. White and Yellow arrowheads denote clone and Control respectively. **(G-I)** Ecdysone pathway depleted follicle cells do not express the centripetal marker *BB127-Lacz*. Clonal cells are marked by GFP, DAPI in cyan and *BB-127* in magenta. **(J)** Plot of cell area of Indicated genotypes. Overexpression of Eya does not rescue the EcRB1^DN^ defect. Error bars indicate SEM. ns = not significant, in Students’ t-test

**Figure S3:**
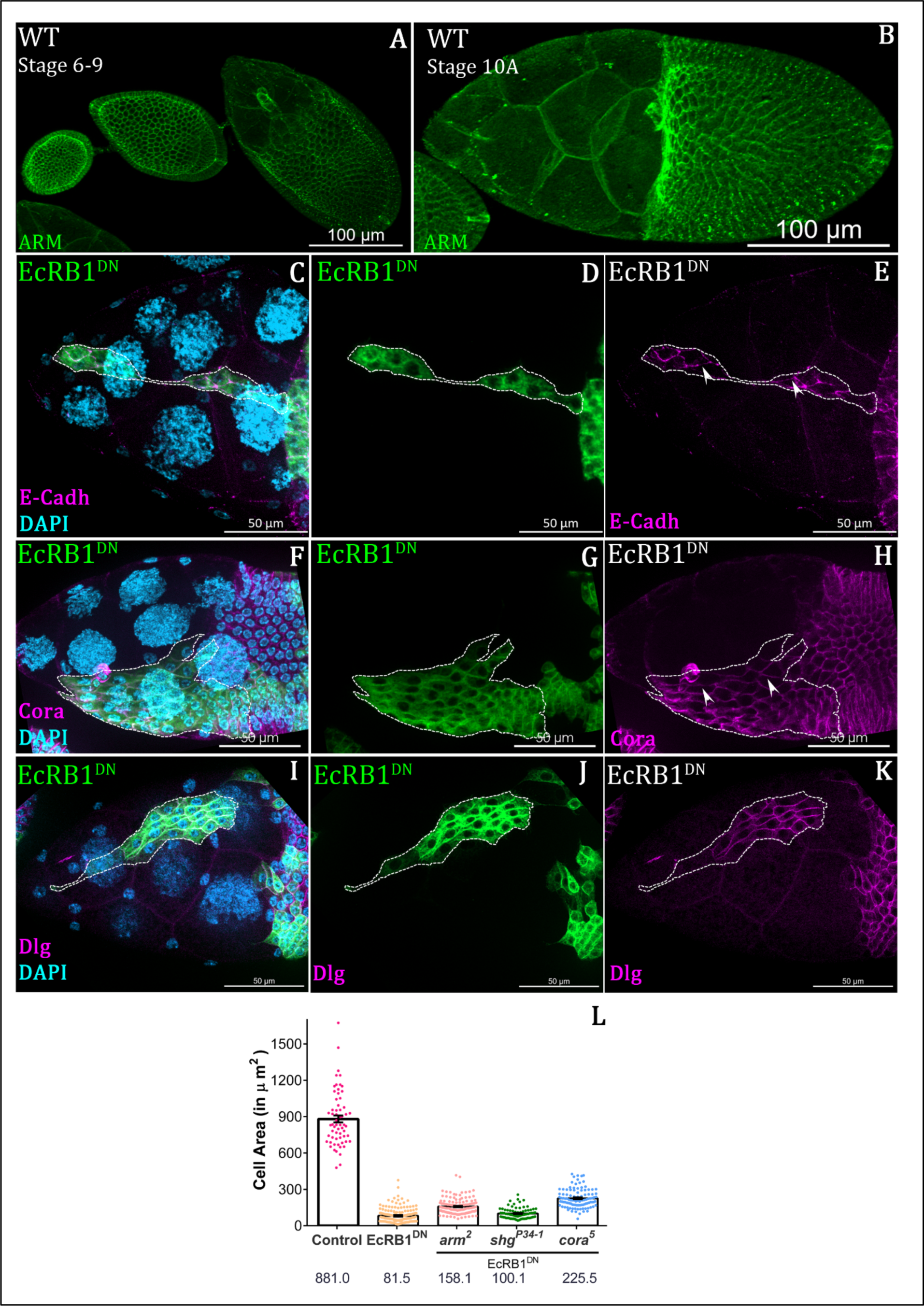
Ecdysone Pathway depleted cells affect the localization of E-Cadh and Coracle. **(A-B)** Expression pattern of Armadillo in wild type Condition at stage 6-9 **(A)** and at stage 10**(B).** Armadillo in green. **(C-H)** Cell junction proteins are accumulated in EcRB1^DN^ clonal population. Staining of E-Cadh **(C-E)**, Coracle **(F-H)**, Dlg **(I-K)**. DAPI in cyan, Clones in green, E-Cadh, Coracle and Dlg in magenta. **(L)** Plot of cell area in the background of indicated genotype. Removal of adherens junction protein does not rescue the defect of EcRB1^DN^. Clonal cells are marked by GFP, DAPI in cyan and E-Cadh or Coracle in magenta. Accumulation of cell junction proteins(E-Cadh and Cora) marked by white arrowhead.

**Figure S4:**
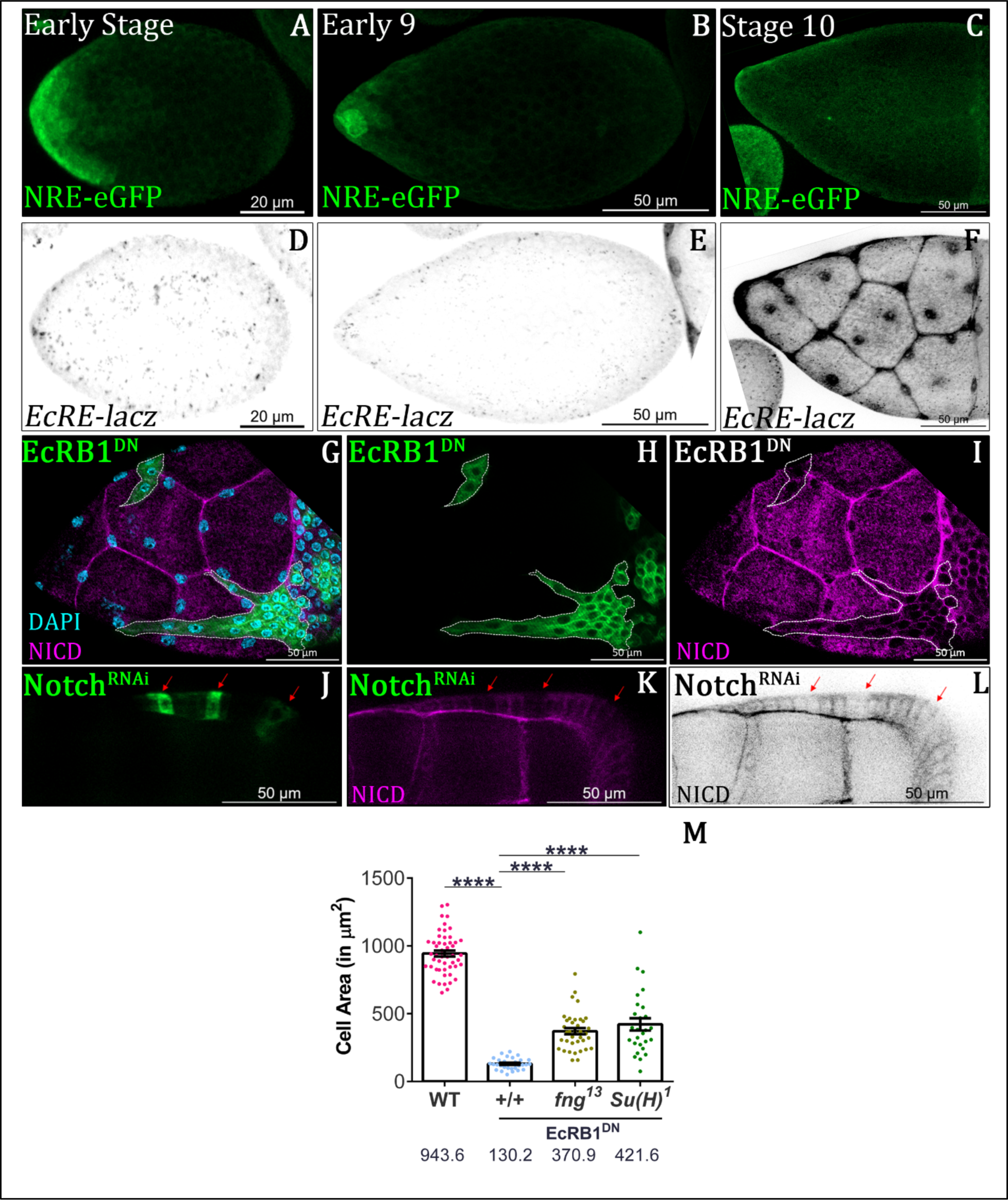
Ecdysone regulates notch in anterior follicle cells. **(A-F)** Stage wise activity of Notch and Ecdysone Pathway. Expression of Notch and Ecdysone pathway are temporally exclusive. NRE-eGFp in green and *EcRE-lacZ* in LUT. **(G-I)** EcRB1^DN^ overexpressed cells exhibit lower level of NICD with respect of neighboring Control cells. Clonal cells are marked by GFP, DAPI in cyan and NICD in magenta. **(J-L)** Level of NICD level of in Notch^RNAi^ background. Clonal cells are marked by GFP, and NICD in magenta. **(M)** Quantification of indicated genotypes. Introduction of heterozygote of *fng^13^* and *Su(H)^1^* in EcRB1^DN^ rescues the stretching defect of EcRB1^DN^. Error bars indicate SEM. **** indicates p-value < 0.0001 in Students’ t-test.

**Figure S5:**
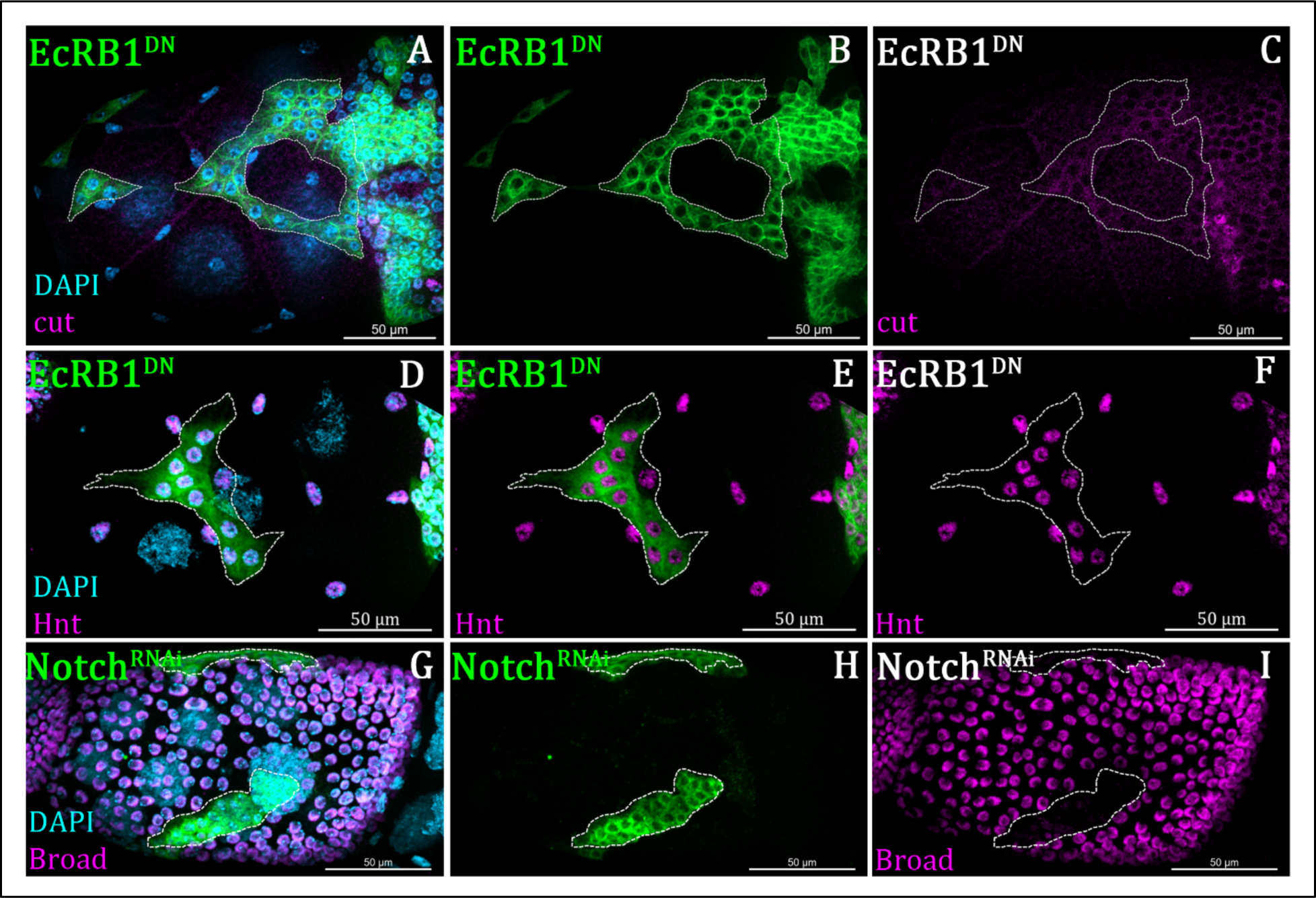
Notch positively regulates Broad. **(A-F)** Ecdysone perturbed cells (in green) do not affect Hindsight **(A-C)** and cut **(D-F)** expression pattern. DAPI in cyan and hnt or cut in magenta. **(G-I)** Reduction of Broad expression in Notch^RNAi^ clones at stage 8. Clones are marked by Flipout mediated mCD8GFP. DAPI in cyan and Broad in magenta.

**Figure S6:**
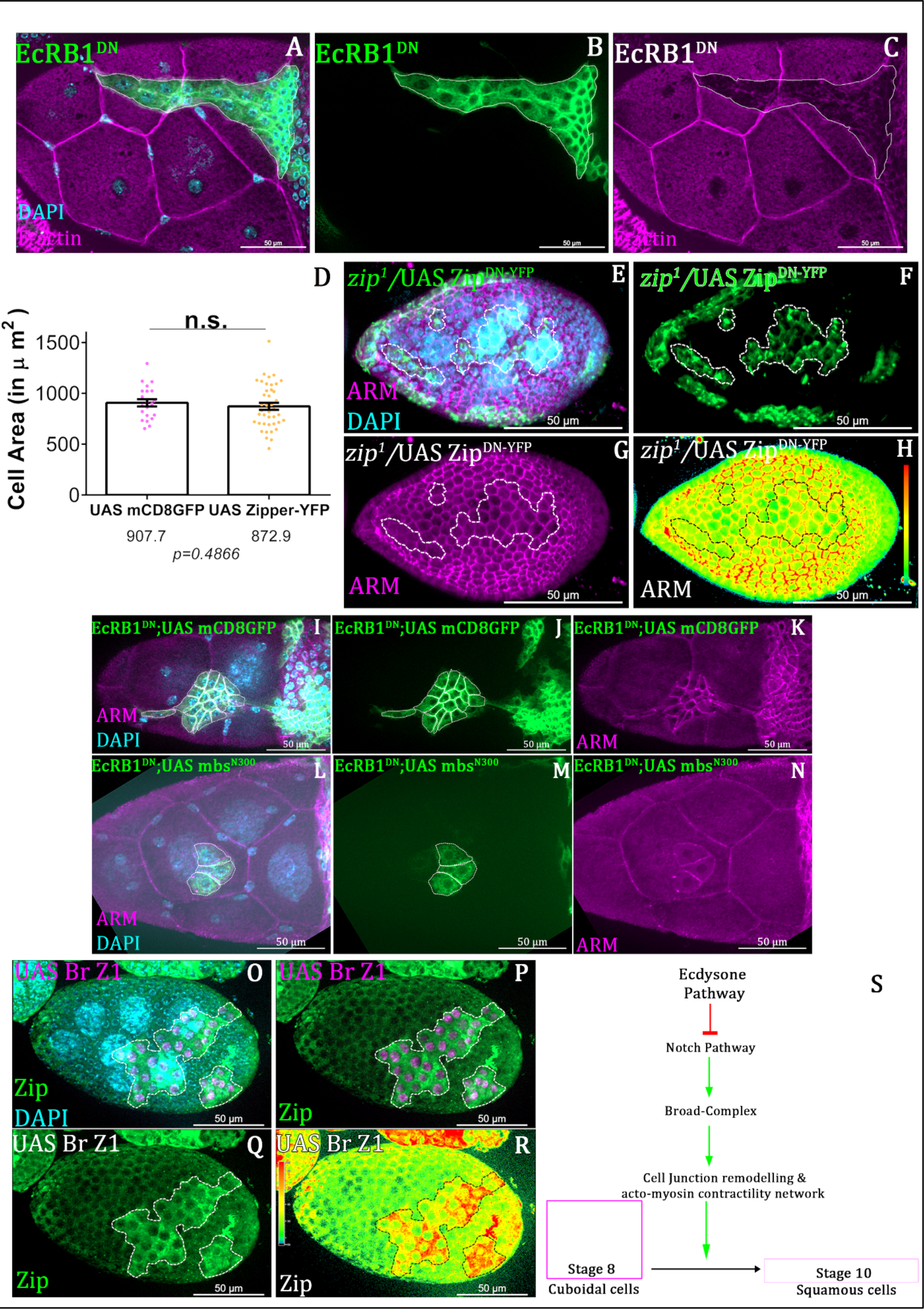
EcR regulates acto-cytoskeleton network. **(A-C)** F-actin levels is reduced in EcRB1^DN^ overexpressed cells. Clones are marked by Flipout mediated mCD8GFP. DAPI in cyan and F-actin in magenta. **(D)** Quantification of Cell area of indicated genotypes. Over-expression of Zipper-^WT^ does not affect cell area in stage 10 **(E-H)** Comparison of cell area at stage 8 of indicated genotypes. Removal of Zipper activity in tranheterozygotes zipper mutant background induces prcocius flattening. Clonal cells marked by Flipout mediated GFP and black dotted line. DAPI in cyan and Armadillo in magenta. **(I-N)** Constitutively active mbs partially rescues stretching defect of EcRB1^DN^. Clonal population of indicated genotypes marked by mCD8GFP and white dotted outlines. Armadillo in magenta and DAPI in cyan. **(O-R)** Elevated zipper expression in the Broad Overexpressed cells at satge 8. Clonal cells are marked by StingerNLS in magenta, DAPI in cyan and zipper in Green. **(S)** Schematic diagrame how ecdysone pathway regulates epithelial morphogenesis. Error bars indicate SEM. ns = not significant, in Students’ t-test

## Notes

### Competing Interest Statement

The authors have declared no competing interest.

